# Biofunctional coating of synthetic magnetic nanoparticles enables magnetogenetic control of protein functions inside cells

**DOI:** 10.1101/2024.10.31.621314

**Authors:** Marie Kappen, Jakub Gemperle, Emilie Secret, Julia Flesch, Patrick Caswell, Mathieu Coppey, Maxime Dahan, Christine Menager, Domenik Liße, Jacob Piehler

**Affiliations:** Department of Biology/Chemistry and Center for Cellular Nanoanalytics, Osnabrück University, 49076 Germany; Wellcome Trust Centre for Cell-Matrix Research, School of Biological Sciences, Faculty of Biology Medicine and Health, Manchester Academic Health Science Centre, The University of Manchester, Manchester, United Kingdom; Laboratory of Integrative Biology, Institute of Molecular Genetics of the Czech Academy of Sciences, Prague, Czech Republic; Physico-chimie des Électrolytes et Nanosystèmes Interfaciaux, PHENIX, Sorbonne Université, CNRS, F-75005 Paris, France; Laboratoire Physico Chimie Curie, Institut Curie, Paris, France

**Author notes:** Deceased (Maxime Dahan).

## Abstract

Remote control of cellular functions via magnetic forces offers unique opportunities in fundamental research and biomedical application. Intracellular delivery of functionalized magnetic nanoparticles (MNP) provides versatile opportunities to assemble signalling platforms for spatiotemporal control by magnetic forces. Such magnetogenetic application, however, has remained highly challenging due to a lack of MNP providing suitable biological, physicochemical and magnetic properties. Here, we achieved single-step surface coating of synthetic maghemite core nanoparticles with green fluorescent protein fused to the iron binding site of Mms6 from magnetotactic bacteria. We yielded MNP with intracellular stealth properties (syMagIcS), which could be readily biofunctionalized *in situ* and translocated within cells via magnetic field gradients. We successfully exploited syMagIcS for spatiotemporal control of Rac1 signalling at the plasma membrane via its guanine nucleotide exchange factor protein TIAM1 and for spatial control of liquid-liquid phase separation using the intrinsically disordered domain of the protein DDX4.

## Introduction

Regulation of protein functions in time and space is a fundamental principle for coordinating cellular processes.^1-4^ Next to direct control of protein functions by enzymatic modification, regulation is often achieved by changing local protein concentrations via selective translocation to cellular membranes or into phase-separated protein droplets. Since the advent of optogenetics, non-invasive control of protein localization by light has provided valuable tools to explore the spatiotemporal regulation of cellular processes.5-9 More recently, magnetic actuation of protein functions via magnetic nanoparticles (MNPs) has augmented the capabilities for remote-controlling biological functions in cells and organisms.^10, 11^ Such “magnetogenetic” manipulation of protein functions has been achieved via MNP-mediated mechanical actuation, local heating or by modulation of protein concentrations in cells^.12-23^ In contrast to optogenetics, magnetic manipulation techniques are readily compatible with opaque specimens, making them particularly attractive for medical application. However, these techniques require equipping target proteins with magnetically controllable properties by conjugation with suitable magnetic nanoparticles (MNPs).^11, 24^ Design of MNP suitable for magnetic manipulation in cells, however, remains challenging. Ideal MNP need to combine (i) high magnetization, (ii) dedicated surface passivation to minimize toxicity^25^ and recognition by cellular degradation machineries^26, 27^ as well as (iii) biofunctionalization for selective and efficient conjugation with target proteins, ideally by *in situ* capturing within cells. At the same time, robust colloidal properties must be maintained while the total hydrodynamic diameter of fully assembled MNPs should not exceed 50 nm to warrant efficient cellular delivery of the nanoparticles and to maintain unhindered mobility in the cytoplasm^28, 29^.

Here, we developed a new strategy for densely coating synthetic maghemite core particles (MCPs) with the highly stable and biologically indifferent mEGFP β-barrels in a single step. This approach yielded synthetic MNPs with intracellular stealth properties (syMagIcS). For efficient, high-affinity binding to MCP surface, mEGFP was fused to a C-terminal 22 amino acid fragment of the iron oxide binding protein Mms6^30-32^ (amino acid 112-133, Mms6ΔN) from magnetotactic bacteria (Fig. 1A, Supplementary Fig. S1A). Mms6ΔN has been shown to be responsible for the very strong iron oxide binding of Mms6,^33, 34^ enabling direct coating of MCPs with mEGFP (Fig. 1B). This coating strategy significantly simplifies the production of biocompatible magnetic nanoparticles, while simultaneously providing the opportunity for selective *in situ* nanoparticle biofunctionalization via GFP binding proteins (Supplementary Fig. S1B). Remarkably, cytosolic stealth properties of mEGFP coated MCPs were achieved without further chemical surface modifications. We demonstrate intracellular inertness of syMagIcS, ultra-fast targeting to cellular proteins as well as rapid attraction and intracellular relocalization by magnetic field gradients. Such “space-mode” magnetogenetic manipulation enabled remote-controlled activation of the GTPase Rac1 at the plasma membrane as well as spatiotemporal control of liquid-liquid phase separation of DDX4 inside living cells.

**Figure 1.**
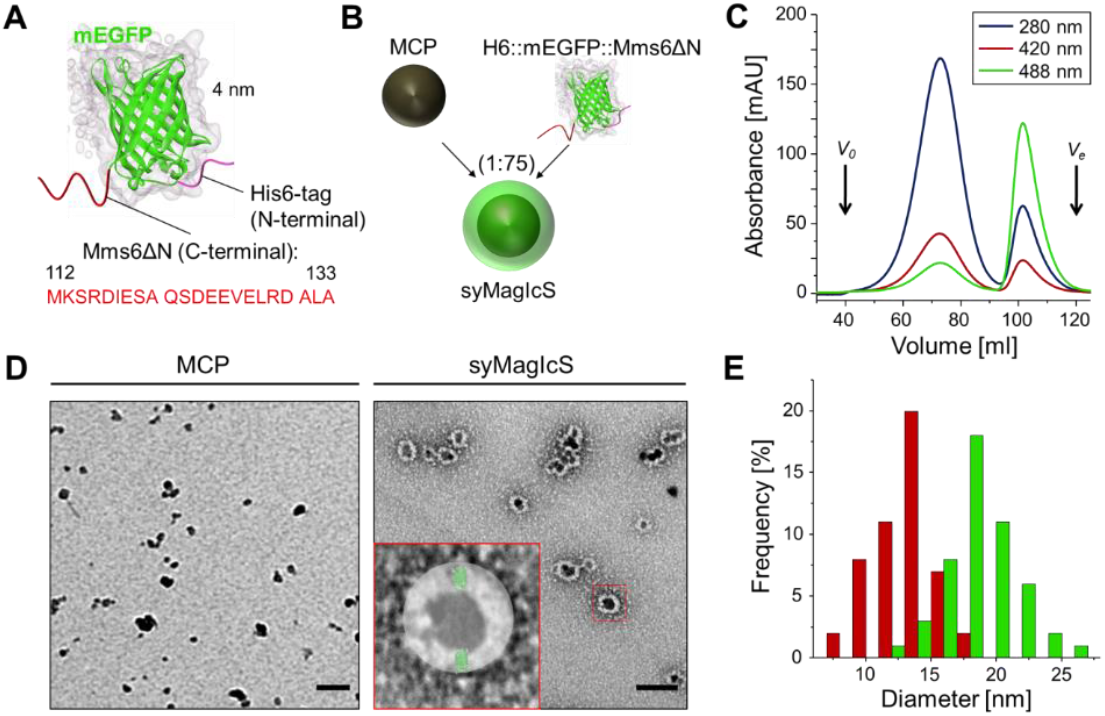
Efficient MCP coating with Mms6ΔN-functionalized GFP. (A-B) Strategy for production of syMagIcS. (A) Schematic model of H6::mEGFP::Mms6ΔN based on a GFP crystal structure (PDB 3K1K). Mms6ΔN is illustrated in red, mEGFP in green and the His_6_-tag in magenta. (B) Coating MCP by mixing a 75-fold excess H6::mEGFP::Mms6ΔN with MCPs at pH 2 in water. (C-F) Physico-chemical characterization of syMagIcS. (C) Analytical size-exclusion chromatography (Sephacryl S500) of the reaction mix detected via absorbance at three different wavelength. Column void volume (*V*_*0*_) and bed volume (*V*_*e*_) are indicated. (D) Transmission electron micrographs of MCPs (left) and syMagIcS (right). Scale bar: 50 nm. (E) Particle size distribution of MCPs (red) and syMagIcS (green) obtained from TEM images; n = 100 particles for each species.

## Results

### Dense coating and high colloidal stability of MCPs achieved with Mms6ΔN-functionalized mEGFP

Recombinant Mms6ΔN tagged mEGFP (H6::mEGFP::Mms6ΔN) was produced in *E. coli* with high yields and purified to homogeneity by immobilized metal affinity chromatography (IMAC) under denaturing conditions to prevent proteolytic degradation followed by size exclusion chromatography (Supplementary Fig. S2). After transferring the protein into pure water, maghemite core particles (MCPs, Supplementary Fig. S3) dispersed in water at pH 2 were added in a protein : MCP ratio of 75:1. Excess of non-bound H6::mEGFP::Mms6ΔN was removed by ultra-filtration. Analytical size-exclusion chromatography in HBS (20 mM HEPES, 150 mM NaCl, pH 8.0) of the reaction mix yielded a monodisperse elution profile of a brownish solution at a volume expected for MCP coated with GFP monolayer (Fig. 1C). While non-coated MCPs precipitate under these conditions, monodisperse elution of those particles indicated effective coating of MCPs with mEGFP. TEM imaging of syMagIcS confirmed complete coating of MCPs by forming a densely packed mEGFP corona around the particles (Fig. 1D). From those images an average physical diameter (*d*_*P*_) of 12.1 ± 2.1 nm for MCPs and 19.6 ± 2.7 nm for syMagIcS, respectively, were observed (Fig. 1E). A similar increase of the hydrodynamic diameter upon MCP coating was observed (Supplementary Fig. S4A). The increased particle size of ∼8 nm is in good agreement with the ∼4 nm height of the mEGFP barrel, corroborating an upright orientation within the MCP coating as expected for the position of the Mms6ΔN peptide fused to the C-terminus (Fig. 1D, inset). Coating with PEGylated H6::mEGFP::Mms6ΔN increased the hydrodynamic diameter to further decreased the aggregation of the syMagIcs (Supplementary Fig. S4B). The ζ-potential of syMagIcS was found to be -25 mV in HEPES buffer pH 7.5, in line with the negatively charged mEGFP under these conditions. Estimation of functionalization stoichiometry yielded an average number of 66 mEGFP molecules per MCP (Supplementary Fig. S5), in line with dense monolayer coverage of the ∼ 430 nm^2^ surface of these particles.

### syMagIcS exhibit high biocompatibility and negligible cytotoxicity

The long-term stability of syMagIcS was probed in minimum essential medium containing 10% fetal bovine serum (MEM^++^) pH 7.5, HEPES buffer pH 7.5 (HB) or Dulbecco’s phosphate-buffered saline (DPBS) (Supplementary Fig. S6). In MEM^++^, particles were stable for several days without significant loss. By contrast, in HB no significant loss of particles were observed over the observation time of 14 days demonstrating tight binding of H6::mEGFP::Mms6ΔN under these conditions. Lower particle stability in culture medium might be explained by the multivalent binding of Mms6ΔN to the maghemite surface^,35^ which can be more readily reversed by competing molecules (e.g. amino acids) in the culture medium resulting particle precipitation. In DPBS a low stability was observed, within one hour most of the particles were lost. However, after crosslinking the GFP shell on the MCP surface by treatment with paraformaldehyde (4% PFA in HB), syMagIcS showed comparable stability in PDBS as in HB.

Since the long-term replacement of the coating protein on the particle surface in biological fluids might yield cytotoxic effects, cell proliferation during three days of culturing was quantified by automated imaging. Electroporation yielded robust and reproducible delivery of syMagIcS to more than > 90 % of cells (Supplementary Fig. S7A). Cytoplasmic syMagIcS had no effect on cell viability suggesting the low cytotoxicity of these nanoparticles (Supplementary Fig. S7B). Only at very high syMagIcS concentrations (at bulk concentrations of ∼1 μM), a minor decrease in the proliferation rate of Hela cells was observed (Supplementary Fig. S7C). In A2780 cells, even the maximum syMagIcS concentrations above 1 μM (Supplementary Fig. S7D) did not cause any decrease in proliferation. Overall, these results highlight that the dense coating with mEGFP yielded highly biocompatible magnetic nanocarriers with very low cytotoxicity.

### syMagIcS are freely dispersed in the cytoplasm and efficiently conjugated with target proteins *in situ*

Recognition by cellular degradation machineries as well as unspecific interaction with large biomolecules or organelles is a key challenge for the application of colloidal nanoparticles inside living cells. Thus, uptake into autophagosomes at a timescale of a few seconds was even found for protein-based nanoparticles.^27^ We therefore investigated the colloidal stability of syMagIcS in the cytoplasm of HeLa cells. Delivery was robustly accomplished by microinjection yielding intracellular syMagIcS concentrations up to 35 nM as quantified by confocal microscopy (Supplementary Fig. S8). Upon microinjection, particles were homogenously distributed in the cytoplasm. Over a period of 30 min, neither particle clustering nor unspecific interaction with the intracellular environment was observed demonstrating advanced intracellular stability (Fig. 2A and Supplementary Movie S1). Thus, the highly dense coating of MCPs with H6::mEGFP::Mms6ΔN was sufficient to yield intracellular stealth properties without the need of additional chemical surface modifications.

**Figure 2.**
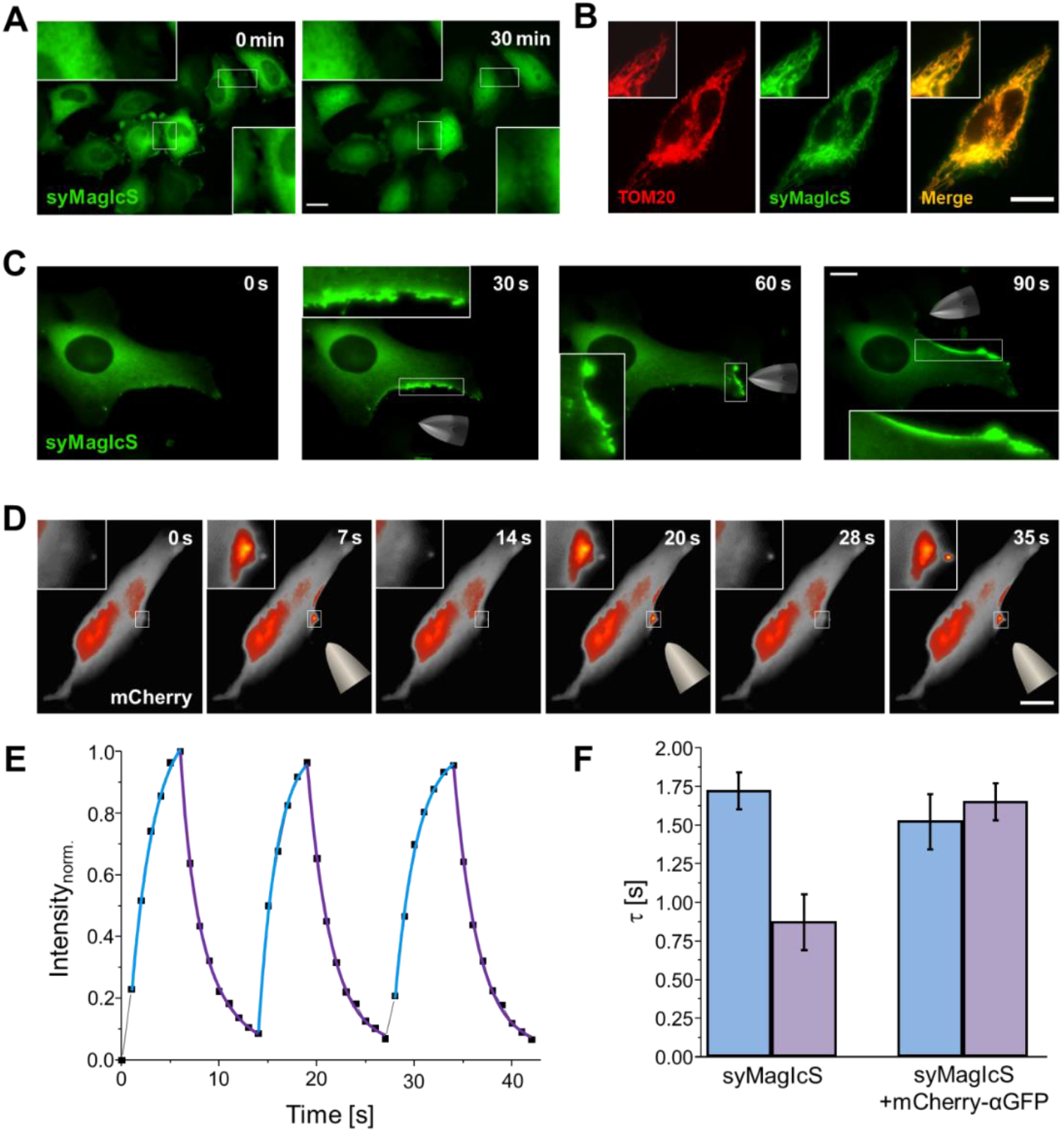
Efficient targeting and magnetic manipulation of syMagIcS inside living cells. (A, B) Distribution of syMagIcS in the cytoplasm of wt HeLa cells (A) and in HeLa cells expressing TOM20::mCherry::αGFP (B). (C) Magnetic manipulation of syMagIcS inside living cells. A magnetic tip was approached to a HeLa cell loaded with syMagIcS (green) from different directions. (D-F) Magnetogenetic manipulation of proteins inside living cells. (D) syMagIcS was injected into HeLa cells expressing mCherry::αGFP and repeatedly attracted and released with a micro-magnet positioned at a ∼30 μm distance to the plasma membrane. Only mCherry fluorescence is shown to demonstrate magnetic manipulation of a cytosolic protein. Scale bars: 30 μm in all images. (E) Attraction and release kinetics of mCherry-functionalized syMagIcS determined from the changes in fluorescence intensity in the area highlighted by the white square. Time constants were determined by fitting an exponential curve (blue, purple). (F) Comparison of the manipulation time constants of attraction and release observed for syMagIcS in the absence and presence of mCherry::αGFP.

Capturing biomolecules to the surface of MNP opens powerful possibilities to assemble active signalling platforms *in situ*. In the case of syMagIcS, we exploited for this purpose the nanobody “enhancer” (αGFP), which binds mEGFP rapidly and with sub-nanomolar affinity, while its binds only very weakly to mECFP.^36^ αGFP has been previously shown to be compatible with intracellular application,^37^ and therefore was genetically fused to intracellular target proteins for efficient, site-specific capturing surface of syMagIcS. *In vitro* binding experiments by solid phase detection confirmed specific recognition of sysMagIcS by αGFP even after fixing the mEGFP coating with PFA (Supplementary Fig. S9). For demonstrating *in cellulo* targeting, TOM20 fused to mCherry and αGFP (TOM20::mCherry::αGFP), which is anchored in the mitochondrial outer membrane with the αGFP facing the cytoplasm, was transiently expressed in HeLa cells. Upon microinjection of syMagIcS, rapid binding to mitochondria was observed on a timescale of a few seconds, yielding highly homogenous co-localization of mEGFP (syMagIcS) and mCherry (TOM20) fluorescence (Fig. 2B). Microinjection of sub-stoichiometric syMagIcS quantities yielded binding to TOM20::mCherry::αGFP only surrounding the micro-injection point (Supplementary Fig. S10A), highlighting the ultra-fast and highly specific *in situ* conjugation of syMagIcS with target proteins possible using this approach. Similarly, efficient targeting of syMagIcS to recycling endosomes was achieved upon microinjection into expression of A2780 cells stably expressing αGFP::mCherry::Rab11A (Supplementary Fig. S11A). Likewise, syMagIcS delivered by electroporation were efficient targeted to Rab11A (Supplementary Fig. S11B).

### Rapid and reversible control of syMagIcS in the cytoplasm by magnetic forces

Fulfilling the key-prerequisites for robust and specific intracellular application of syMagIcS, we next investigated magnetic manipulation of freely diffusing particles and magnetogenetic manipulation of proteins inside HeLa cells. To this end, we employed a home-built micro-magnet assembled of commercially available parts, which was mounted via an electronic micromanipulator to an epifluorescence microscope (Supplementary Fig. S12). At first, we tested the response of freely diffusing syMagIcS after microinjection into HeLa cells. Strikingly, when approaching the micro-magnet at ∼20 μm distance to the cell membrane (Fig. 2C and Supplementary Movie S2), rapid attraction of syMagIcS could be observed at the cytoplasmic side close to the micro-magnet within a few seconds. Upon removing the micro-magnet, particles diffused back toward the cell interior and could be readily manipulated to different subcellular positions within the cell. Since syMagIcS could be repeatedly attracted and released inside living cells (Supplementary Fig. S13), evaluation of attraction and relaxation time constant yielded typical values of *τ*_*A*_ = 1.7 ± 0.1 s and *τ*_*R*_ = 1.0 ± 0.2 s when the micromagnet was at 45 μm distance to the plasma membrane, demonstrating that fast spatial-temporal magnetic control of syMagIcS was possible (Supplementary Fig. S13 and Supplementary Movie S3). The magnetic force exerted on a particle at 48 μm to 152 μm distance to the micro-magnet was in the Femto-Newton range, as determined using the Boltzmann law from the steady-state profile of syMagIcS distribution inside living cells (Supplementary Fig. S14).

To demonstrate magnetogenetic manipulation, *i*.*e*. spatial control of a genetically encoded target protein expressed in a cell, we microinjected syMagIcS into HeLa cells transiently overexpressing αGFP fused to mCherry (mCherry::αGFP). Upon placing the magnetic tip at ∼30 μm distance to the plasma membrane, mCherry::αGFP rapidly accumulated in proximity to the tip (Fig. 2D and Supplementary Movie S4). Evaluation of the attraction and relaxation kinetics yielded time constants *τ*_*A*_ = 1.7 ± 0.2 s and *τ*_*R*_ = 1.5 ± 0.1 s, *i*.*e*. similar as for syMagIcS only (Fig. 2E, F). The reduced relaxation time constants observed in the presence of mCherry::αGFP could be ascribed to the increased hydrodynamic diameter upon binding to the syMagIcS surface. Rapidly changing the tip position within a time period of 6 s yielded a velocity of ∼5 μm/s for moving mCherry-functionalized syMagIcS within the cytoplasm (Supplementary Fig. S13 and Supplementary Movie S5). These results demonstrate highly robust, reversible and fast relocalization of proteins in the cytosplasm by magnetic of syMagIcS.

### Spatially controlled activation of the GTPase Rac1 by magnetogenetic manipulation

To demonstrate the capability of magnetogenetic activation of biological processes, we chose the Rho family GTPase Rac1 as a model system. Rho family GTPases regulate and coordinate cytoskeleton remodelling by inducing polymerisation of actin,^38, 39^ with Rac1 being mostly involved in the formation of protrusions.^40, 41^ Here, we explored the activation of Rac1 by translocating its guanine nucleotide exchange factor (GEF) TIAM1 via magnetic forces to the plasma membrane (Fig. 3A). To this end, syMagIcS was employed as a controllable nanocarrier to control subcellular localization and thus activity of TIAM1. By doing so, we first investigated the capability to recruit Rac1 to syMagIcS functionalized with the catalytically active fragment of TIAM1 comprising the DH and PH domains (TIAM1-DHPH, amino acid 1033-1406). This fragment lacks the ability of the full-length TIAM1 to localize to the plasma membrane. To monitor the recruitment of Rac1, syMagIcS were injected into COS7 cells expressing TIAM1-DHPH fused to mCherry and αGFP (TIAM1-DHPH::mCherry::αGFP) together with Rac1 fused to the HaloTag (HaloTag::Rac1), which was stained with silicon rhodamine (SiR). Recruitment of Rac1 by syMagIcS, which was *in situ* functionalized with TIAM1-DHPH (TIAM-syMagIcS), was observed upon translocation to the plasma membrane by means of magnetic forces (Fig. 3B). On a timescale of a few minutes, Rac1 substantially accumulated at the site where TIAM-syMagIcS was dragged to the plasma membrane (Fig. 3B and Supplementary Movie S6). TIAM-specific magnetogenetic Rac1 recruitment was robust and could be reproducibly observed in 90% of executed experiments. In control experiments without TIAM captured to syMagIcS, Rac1 recruitment could not be discerned (Supplementary Fig. S16A).

**Figure 3.**
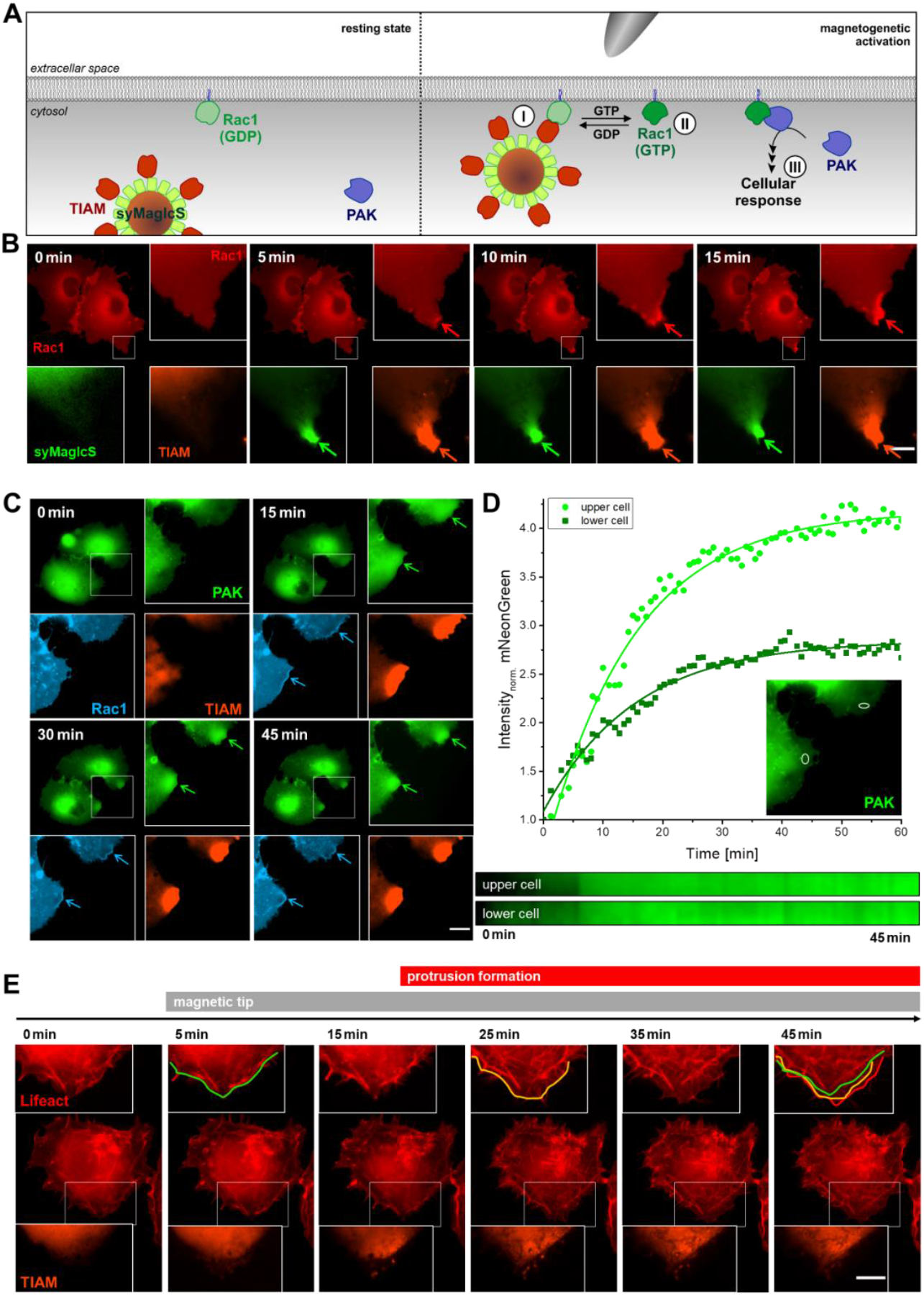
Magnetogenetic activation of Rac1. (A) Cartoon depicting the strategy for magnetogenetic control of Rac1: translocating the GEF protein TIAM captured to syMagIcS to the plasma membrane, interaction with Rac1-GDP (step I) promotes GDP-GTP exchange (step II). Rac1-GTP recruits the effector protein PAK to initiate downstream activity leading to formation of cell protrusions (step III). (B) Recruitment of Rac1 by TIAM-functionalized syMagIcS translocated to the plasma membrane via magnetic forces (step I): syMagIcS (green) was injected into cells expressing HaloTag::Rac1 labeled with SiR (red) and TIAM1-DHPH::mCherry::αGFP (orange). Time-lapse imaging during magnetic manipulation showing Rac1 distribution in the entire cell and zoom into the zone enriched in syMagIcS for all three channels. (C) Magnetogenetic control of Rac1 GDP-GTP exchange (step II) detected by monitoring recruitment of PAK-CRIB: syMagIcS (colorless) was injected into cells expressing TIAM1-DHPH::mCherry::αGFP (orange) and mNG::PAK-CRIB::P2A::Rac1::mTFP1::CAAX (green/cyan). Time-lapse imaging during magnetic manipulation showing PAK-CRIB distribution and zoom into the zone enriched in syMagIcS for all three channels. The kinetics of PAK-CRIB recruitment is shown by the intensity increase of mNeonGreen (top) and corresponding kymographs (bottom). (D) Kinetics of mNG::PAK-CRIB recruitment within the two white ROIs indicated in the inset and corresponding kymographs (bottom). (E) Magnetogenetic activation of Rac1 downstream signal responses by monitoring protrusion formation (step III): syMagIcS was injected into cells expressing TIAM1-DHPH::mCherry::αGFP (orange) and Lifeact::iRFP (red) for staining f-actin. Time-lapse imaging during magnetic manipulation showing the actin cytoskeletal structure and zoom into the zone enriched in TIAM for both channels. Scale bars: 10 μm in all images.

To test magnetogenetic activation of Rac1 GDP-GTP exchange (Fig. 3A, step II), recruitment of the Rac1-GTP effector protein PAK was probed. To this end, the Rac1 binding domain CRIB of PAK (PAK-CRIB) fused to mNeonGreen (mNG) was co-expressed with Rac1 fused to mTFP1 using the P2A peptide linker to ensure similar protein levels (mNG::PAK-CRIB::P2A::Rac1::mTFP1::CAAX) and TIAM1-DHPH::mCherry::αGFP for syMagIcS functionalization. To avoid interfering fluorescence, non-fluorescent H6::mXFP::Mms6ΔN (mEGFP Y67F) was used for coating MCP. Upon translocation of TIAM-syMagIcS to the plasma membrane, recruitment of Rac1 was observed, which was followed by local enrichment of PAK (Fig. 3C, D and Supplementary Movie S7). By contrast, in control experiments without TIAM1-DHPH, neither Rac1 nor PAK recruitment were observed (Supplementary Fig. S16B). Strikingly, downstream cellular responses of locally activated Rac1 could also be detected (Fig 3A, step III). To this end, syMagIcS were injected into COS7 cells co-expressing TIAM1-DHPH::mCherry::αGFP and Lifeact fused to iRFP (Lifeact::iRFP). Upon attraction with the magnetic tip, we observed the local enhancement of actin polymerization and protrusion formation (Fig. 3E and Supplementary Movie S8). Interestingly, we observed stronger responses in regions with lower particle densities (Supplementary Movie S9 and Fig. S15), pointing towards some autoinhibitory effect of TIAM-syMagIcS. In general, weaker cellular responses were observed compared to optogenetic activation of Rac1,^42^ which may be related to this autoinhibitory effects.

### DDX4-functionalized syMagIcS enables spatial control of protein droplet formation and localization

Liquid-liquid phase separation (LLPS) mediated by multivalent interactions of intrinsically disordered protein regions (IDRs) has recently emerged as a key principle in the spatiotemporal organization of fundamental cellular functions such as transcription and translation, cell division and signalling^.43-48^ Concomitantly, LLPS has been implicated in several pathophysiological processes involved in complex diseases such as cancer and neurodegeneration.^49-51^ Providing the ability to remote-control concentrations of cellular proteins in time and space, we here explored magnetogenetic manipulation of LLPS. As a model system, the IDR of DDX4, which have been reported to drive LLPS of DDX4,^52^ was fused to mCherry and αGFP (DDX4::mCherry::αGFP) for *in situ* capturing by syMagIcS (Fig. 4A).

**Figure 4.**
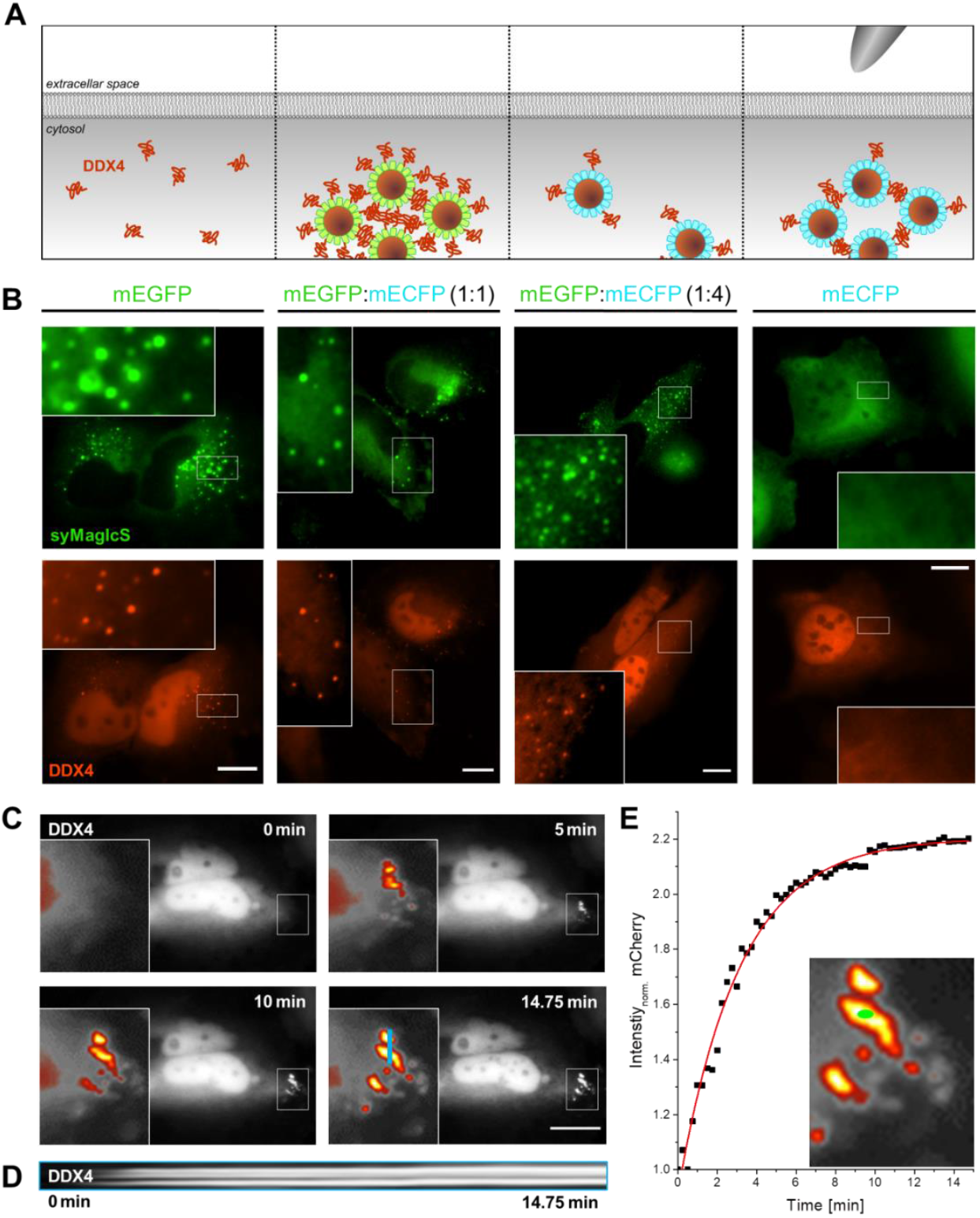
Spatiotemporal control of protein condensation. (A) Strategy: syMagIcS is *in situ* functionalized with the IDR of DDX4 and phase-separation is induced by locally increasing the concentration via magnetic forces. The multivalency induced by DDX4 binding to the syMagIcS surface is reduced by including mECFP into the MCP coating, which binds αGFP with ∼1000-fold reduced affinity (*K*_D_: 450 nM). (B) Spontaneous phase separation upon injection of syMagIcS coated with different ratios of mEGFP and mECFP into HeLa cells expressing DDX4::mCherry::αGFP. (C) Magnetogenetic controlled phase-separation inside living cells. syMagIcS coated with mECFP were microinjected into HeLa cells overexpressing IDR::mCherry::anti-GFP nanobody. Time-lapse imaging of phase-separation upon applying magnetic forces. (D) Kymograph along the cyan line as indicated in panel C. (E) Kinetics of DDX4 phase separation within the green ROI shown in the inset. Scale bars: 20 μm in all images.

After micro-injection of syMagIcS into HeLa cells expressing DDX4::mCherry::αGFP, spontaneous LLPS was observed on a timescale of seconds without magnetic stimuli (Fig. 4B), which probably was driven by an increased DDX4 multivalency upon binding to the syMagIcS surface. We therefore reduced the effective surface multivalency by MCP coating with mECFP, which binds αGFP with a ∼1000-fold decreased binding affinity (*K*_D_: 450 nM). While syMagIcS coated with mixtures of mEGFP and mECFP up to a ratio of 1:4 still induced spontaneous LLPS, injection of MCPs solely coated with mECFP did not change the phase behaviour inside cells (Fig. 4B). By contrast, subcellular magnetic attraction of mECFP-coated syMagIcS resulted in a spinodal-like phase separation within minutes (Fig. 4C and Supplementary Movie 10). In some cases, we also could observe nucleation of DDX4, which may be ascribed to varying magnetic field gradients or protein and/or syMagIcS concentrations. While additional studies are needed to realize the full potential of this approach, these results demonstrate the feasibility of magnetogenetics for spatiotemporally controlling phase separation inside living cells.

## Discussion

These cellular applications establish syMagIcS as a novel, versatile tool to control protein function in time and space via magnetic forces. Major asset of syMagIcS is its high compatibility with the intracellular environment yielding low cytotoxicity and high colloidal stability in the cytoplasm. Compared to magneto-ferritin based MNP,^37^ syMagIcS showed much enhanced responsiveness to magnetic field gradients. Production of syMagIcS is very robust and simple, solely requiring mixing of MCP with the coat protein, which can be readily produced in large quantities using a bacterial expression system. Intracellular delivery of syMagIcS in large quantities was readily achieved by microinjection and by electroporation. Given the high colloidal stability, we expect similar compatibility with other methods such as bead loading or micropinocytosis, which have been successfully employed for nanoparticles with similar size and physicochemical properties. The inbuilt surface functionalization implemented with the biocompatible surface coating not only allows efficient *in situ* capturing of target proteins via αGFP, but also to control the degree of functionalization using corresponding GFP mutants. Exploiting these features, we successfully demonstrate spatiotemporal control of G-protein signalling at the plasma membrane by “space-mode” magnetogenetics. By further-more adapting this technology for initiating protein condensation inside living cells, we here provide a unique toolbox for non-invasive manipulation of cellular behaviour in complex environment. While these biological systems have been successfully controlled via optogenetics,^53^ magnetogenetic manipulations offers the unique opportunity to position signalling platforms inside cells. This feature opens novel avenues to unravel the emerging intricate interplay between LLSP and membrane organization, establishing space-mode magnetogenetics as an important, complementary approach. Moreover, exciting biomedical applications, e.g. in regenerative medicine can be envisaged.

## Acknowledgements

We thank A. Budke-Gieseking, G. Hikade, H. Kenneweg and W. Kohl for technical support, and R. Kurre (Integrated Bioimaging Facility Osnabrück) for support with fluorescence microscopy. This project has been funded the European Union’s Horizon 2020 Research and Innovation Programme under grant agreement no. 686841 (MAGNEURON) to M.C., C. M. and J.P. and no. 836212 (EN-DOPOS) to J.G. P.C. is supported by the MRC (MR/R009376/1).

## Author contributions

D.L. and J.P. conceived the project. M.K., D.L., J.G, J.F. performed experiments. E.S. provided materials. P.C., M.C., C.M. and J.P. supervised the project and acquired funding. D.L., M.K. and J.P. wrote the manuscript with input from all other authors.

## Materials and Methods

### Plasmids

| Plasmid | Vector | Insert |
| --- | --- | --- |
| H6::mEGFP::Mms6ΔN | pet21a | 6xHis-tag, linker (3 amino acids), mEGFP, linker (7 amino acids), Mms6ΔN (Mms6, amino acid 112-133, Q2W8R5) |
| H6::mXFP::Mms6ΔN (mEGFP Y67F) | pet21a | 6xHis-tag, linker (3 amino acids), mXFP, linker (7 amino acids), Mms6ΔN (Mms6, amino acid 112-133, Q2W8R5) |
| H6::mCFP::Mms6ΔN (mEGFP H164N) | pet21a | 6xHis-tag, linker (3 amino acids), mCFP, linker (7 amino acids), Mms6ΔN (Mms6, amino acid 112-133, Q2W8R5) |
| Tom20::mCherry::αGFP | pSems | human Tom20 (Q15388), linker (16 amino acids), mCherry, linker (7 amino acids), αGFP |
| TIAM1-DHPH::mCherry::αGFP | pSems | TIAM1-DHPH (mouse TIAM1, amino acid 1033-1406, Q60610), linker (24 amino acids), mCherry, linker (7 amino acids), αGFP |
| αGFP::mCherry::SOScat | pSems | αGFP, linker (3 amino acids), mCherry, linker (8 amino acids), SOScat (human SOS1, amino acid 564-1049, Q07889) |
| HaloTag::Rac1 | pSems | HaloTag, linker (17 amino acids), human Rac1 (P63000) |
| mNeonGreen::PAK-CRIB::P2A::Rac1::mTFP1::CAAX Box | pSems | mNeonGreen, linker (1 amino acids), PAK-CRIB (human PAK1, amino acid 68-150, Q13153), P2A, human Rac1 (amino acid 1-176, P63000), linker (2 amino acids), mTFP1, linker (2 amino acids), CAAX Box |
| mNeonGreen::hRas::EVlinker::RafRBD::mTFP1::CAAX (Ras-Raichu biosensor) | pSems | mNeonGreen, linker (2 amino acids), human hRas (amino acid 1-172, P01112), EVlinker (119 amino acids), Ras binding domain of Raf (human Raf1, amino acid 51-131, P04049), linker (2 amino acids), mTFP1, linker (2 amino acids), CAAX Box |
| Lifeact::iRFP | pIRES | Lifeact, linker, iRFP |
| DDX4::mCherry::αGFP | pSems | DDX4ΔC (human DDX4, amino acid 1-236, Q9NQI0), linker (28 amino acids), mCherry, linker (7 amino acids), αGFP (nanobody enhancer) |
| αGFP::mCherry::Rab11A | pCDH (lentiviral) | αGFP, linker (6 amino acids), mCherry, linker (35 amino acids), human Rab11A (P62491) |
| αGFP::mCherry | pSems | mCherry, linker (7 amino acids), αGFP (nanobody enhancer) |

#### Cloning, expression and purification of mEGFP-Mms6 fusion proteins

The GFP-Mms6 fusion proteins for bacterial expression were cloned into the vector pET21a. The iron-binding fragment of Mms6 comprising the C-terminal 22 amino acid residues (MKSRDIESA QSDEEVELRD ALA) of Mms6 (Mms6ΔN) was genetically fused to the C-terminus of monomeric enhanced GFP (mEGFP), the non-fluorescent mutant mEGFP Y66F (mXFP), or monomeric enhanced CFP (mECFP), containing 7 amino acids as linker by cassette cloning. For purification, a 6xHis-tag was fused to the N-terminus of the respective GFP variant. For bacterial expression of pET21a-H6::mEGFP::Mms6ΔN, pET21a-H6::mXFP::Mms6ΔN, or pET21a-H6::mECFP::Mms6ΔN in *E. coli* BL21-CodonPlus-RIL, the cDNA of the fusion protein was cloned into pET21a (Novagen). Cells were grown at 37 °C, expression was induced with 0.5 mM IPTG (isopropyl β-D-1-thiogalactopyranoside, Thermo Fisher Scientific) at OD_600_ = 0.6-0.8 and subsequently cultured at 16 °C overnight. Cells were harvested by centrifugation at 6500 g for 10 min and subsequently suspended in lysis buffer (50 mM HEPES, 150 mM NaCl, 8 M urea, pH 7.5-8.0). The cell lysate was lysed by tip sonication (4x 3 min; 50% duty cycle, Sonifier 250, Branson) and insoluble material was pelleted by centrifugation at 20000 g for 30 min. The protein was then purified by immobilized metal affinity chromatography and size exclusion chromatography performed in an FPLC system (Äkta Explorer, GE Healthcare). The supernatant was first loaded on a 5 mL HiTrap chelating HP column loaded with nickel (II) chloride and equilibrated in the 20 mM HEPES, 150 mM NaCl, pH 8.0 (HBS). The protein was eluted with HBS complemented with 500 mM imidazole, pH 8.0 using a linear gradient covering 10× the bed volume. The eluted protein was loaded on a SEC column (HiLoad 16/60 Superdex 75, prep grade or HiLoad 26/60 Superdex 75, prep grade) equilibrated in HBS. Protein integrity and purity were confirmed by 12 % SDS-PAGE (MW: 31.2 kDa). H6::mXFP::Mms6ΔN was PEGylated by incubating a 30-fold excess of PEG NHS ester (CH_3_O-PEG-NHCO-C_2_H_4_-CONHS, 2000 Da, Rapp Polymere) for >1 h followed by SEC (HiLoad 16/60 Superdex 200 prep grade, HiLoad 26/60 Superdex 200 prep grade, or Superdex 200 10/300 GL; equilibrated in 20 mM HEPES, 150 mM NaCl, pH 8.0). In the following the purified protein is called coating protein.

#### Synthesis of maghemite MCP

Maghemite nanoparticles were synthesized by an inverse co-precipitation method. The acidic iron (II) and iron (III) ions solution (248.5 g of FeCl_2_,4H_2_O, 100 mL of HCl 37%, 500 mL of DI water, 587 mL of FeCl_3_ 27%) was added dropwise over 30 h into 4 L of 5 % ammonia in water under agitation. After rinsing the obtained Fe_3_O_4_ nanoparticles with water, they were redispersed in 2 L of nitric acid (9.5 %). These nanoparticles were then oxidized into γ-Fe_2_O_3_ nanoparticles by boiling them with a solution of iron (III) nitrate (323 g of Fe(NO_3_)_3_ in 800 mL of DI water) for 30 min. After washing the nanoparticles with nitric acid once, acetone three times and diethyl ether twice, they were redispersed in 1 L of water. At this step the maghemite nanoparticles are polydispersed in size. To decrease the polydispersity, the nanoparticles were size-sorted by addition of nitric acid which increase the ionic strength of the solution, leading to the flocculation of the larger, thus less stable, nanoparticles. The final nanoparticles were size sorted in two steps: first with 20 mL of nitric acid (68%), then with 5 mL of nitric acid. After each size sorting step, the nanoparticles were rinsed with acetone 3 times and with diethylether twice before being resuspended in DI water. The obtained maghemite nanoparticles were characterized by transmission electron microscopy (TEM) on a JEOL 100CX2 instrument, by dynamic light scattering (DLS) on a Malvern Zetasizer NanoZS instrument and by superconducting quantum interference device (SQUID) magnetometry on a Quantum Design MPMS-XL instrument (Supplementary Figure S3 and S4).

#### Functional coating of maghemite core particles

Prior to MCP coating, H6::mEGFP::Mms6ΔN was transferred into to water pH 7.0 using a buffer exchange column (NAP5, NAP10 or PD10). After sonication (Ultrasound bath, 10 min, 10-15 °C) MCP were added to a 75-fold molar excess of H6::mEGFP::Mms6ΔN and incubated for > 1 h. Excess protein was washed out by ultrafiltration (Amicon UF-Tubes, cut-off: 100 kDa, for V ∼ 1000 μL, 4 min, 4000 g, 20 mM HEPES pH 7.5, RT) until the filtrate was free of coating protein. For optional crosslinking of the coating protein shell, particles were incubated in paraformaldehyde (4% PFA in HB, > 1 h, RT). This was followed by a washing step using ultrafiltration (Amicon UF tubes, cut-off: 100 kDa, ∼ 1000 μL, 4 min, 20 mM HEPES pH 7.5, > 3 times, RT). Directly before use inside living cells, particles were centrifuged to remove aggregates at 600 g for 5 min.

#### Fabrication of magnetic tips

Tips for magnetic manipulation were home-built. For this purpose, an iron string (0.1 mm diameter) was pulled in the flame of a Bunsen burner. The string was pulled slowly, resulting in two sharp extremities of 20 μm diameter, which were used as paramagnetic tips mounted on top of a small permanent magnet of neodymium iron boron N-52 (dimensions 5 mm × 2 mm × 1 mm, magnetic poles axial over the length, HKCM, Eckernförde).

#### Cell culture, transfection, TMRM incubation and microinjection

HeLa and COS7 cells were cultivated at 37 °C, 5% CO_2_ in MEM (PAA) with 1.1 % HEPES (PAA), 1.1% NEA (Biochrom/PAA), 10% fetal bovine serum (Biochrom) (MEM^++^) and seeded on sterilized glass coverslips in 35 mm cell culture dishes. After 6 to 24 hours cells were transfected using Viafect reagent (Promega, Madison, WI, USA, ratio: 1 μg DNA: 3 μL Viafect) according to the protocol. The next day cells were washed twice with DPBS buffer and media was exchanged for fresh MEM++. In case of HaloTag an additional labeling step is needed (wash with preheated DPBS, incubate 20 min at 37 °C with 50 nM SIR-HTL in MEM^++^, wash three times with DPBS, add fresh MEM++) is needed. To increase the brightness of iRFP 25 μM biliverdin was added 16 h before the experiment (wash three times with DPBS, add fresh MEM^++^ before imaging). syMagIcS were microinjected into cells based on published protocols^54^ using a micro manipulation system (InjectMan NI2 and 10 FemtoJet Express, Eppendorf) and a capillary pressure between 15-25 hPa. Injection needles (GB100TF-10, 0.78 x 1.00 x 100 mm, Science Products GmbH) were pulled by a P97 micropipette puller (Sutter Instruments, Novato, CA, USA) using the following parameters: heat: 438; pull: 130; velocity; 38; Time: 105. For microinjection the tip of the needle was slowly approached until it touched the cell membrane and a smooth flow of liquid entering the cell was observed. After injection, waiting for 5-10 minutes allowed the cells to recover.

#### Electroporation and cytotoxicity assays

syMagIcS were electroporated into HeLa or A2780 cells using an Amaxa Nucleofector II Electroporation Machine (Lonza) on program A-023. In detail, cells from one sub-confluent 15 cm plate per condition were trypsinized, washed 1x with PBS and 1x in Opti-MEM and finally resuspended in 100 μL of Opti-MEM media (containing 20 mM HEPES). Just prior to electroporation, syMagIcS were added directly to the cells and transferred to a 0.2 cm cuvette. Immediately after electroporation (program A-023), all cells were transferred to 14 mL of pre-warmed complete media. The cytotoxicity of syMagIcS was investigated by an IncuCyte automated imaging system (Essen BioScience; 3 independent repeats). Briefly, 15,000-35,000 cells (corresponding to approximately 100 μL of electroporated cells transferred to 14 mL complete media: HeLa in DMEM media; A2780 in RPMI media) were seeded onto 96-well plates in at least quadruplicates for each condition and wells corresponding to 40% of cell confluence used for automated analysis. The IncuCyte system was programmed to obtain real-time phase-contrast images (10× objective) of the wells every 30 min for 3 days. The IncuCyte imaging system was then used to automatically calculate the area of each cell-covered zone at each time point up to the point of complete closure from an average of the quadruplicate.

#### Imaging and image analysis

For Imaging cells were grown on 25 mm glass coverslips. Imaging and manipulation were performed at Zeiss Axiovert 200 equipped with 40× water-objective and a Zeiss Axiocam. Confocal laser scanning microscopy images was taken at Olympus cLSM FV-1000 equipped with 40× water-objective. Image analyzing was done with Fiji-ImageJ (NIH, USA) and FRET was analyzed with self-written routines in Matlab (R2017a, The Mathworks Inc, Natick, MA).

#### Transmission Electron Microscopy (TEM) images

syMagIcS was applied onto negatively glow-discharged carbon-coated grids (400 mesh, copper grid) for 1 minute. Excess liquid was removed by blotting with filter paper. The grid was washed twice shortly with distilled water and stained 2 min with 1% uranyl acetate and blotted. Digital micrographs were collected using a Zeiss LEO 912 electron microscope operated at 80 kV equipped with a TRS sharp:eye dual speed 2k-on-axis Digital CCD Camera. Image analyzing was done with Fiji-ImageJ.

#### Interaction analysis by TIRFS-RIF detection

*In vitro* quantification of the interaction of αGFP with syMagIcS were carried out by simultaneous total internal reflectance fluorescence spectroscopy and reflectance interference (TIRFS-RIf) detection using home-built setup described previously^55^. Glass substrates coated with a 325–400 nm silica layer were used as transducers for RIf detection. Transducer slides were plasma cleaned (10 min) and then sandwich-incubated (10 min) with 6 μL poly-L-lysine-graft-poly(ethylene glycol) functionalized with the HaloTag ligand (PLL-PEG-HTL)^56^ (1 mg/mL). The chips subsequently were rinsed in MilliQ, dried with nitrogen and optionally stored at -20°C. Data analysis was performed using OriginPro9.1 (OriginLab) and BIAevaluation 3.1 (Cytiva).

#### Quantification of forces exerted by magnetic manipulation

The applied force on syMagIcS was determined in the cell cytoplasm at the steady-state of attraction. The distance-dependent decrease of the particle intensity *I*(*r*) was fitted exponentially. Assuming that the theoretical profile follows the Boltzmann law:

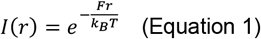

The applied force *F* was determined from the fitted exponential decay (*k*_*B*_: Boltzmann constant; *T*: temperature).

## References

1. Scott, J.D. & Pawson, T. Cell signaling in space and time: where proteins come together and when they’re apart. Science 326, 1220–1224 (2009).

2. Kholodenko, B.N., Hancock, J.F. & Kolch, W. Signalling ballet in space and time. Nature Reviews Molecular Cell Biology 11, 414–426 (2010).

3. Vartak, N. & Bastiaens, P. Spatial cycles in G-protein crowd control. EMBO J 29, 2689–2699 (2010).

4. Dehmelt, L. & Bastiaens, P.I. Spatial organization of intracellular communication: insights from imaging. Nat Rev Mol Cell Biol 11, 440–452 (2010).

5. Guntas, G. et al. Engineering an improved light-induced dimer (iLID) for controlling the localization and activity of signaling proteins. Proceedings of the National Academy of Sciences 112, 112–117 (2015).

6. Rost, B.R., Schneider-Warme, F., Schmitz, D. & Hegemann, P. Optogenetic Tools for Subcellular Applications in Neuroscience. Neuron 96, 572–603 (2017).

7. Toettcher, Jared E., Weiner Orion D. & Lim Wendell A. Using Optogenetics to Interrogate the Dynamic Control of Signal Transmission by the Ras/Erk Module. Cell 155, 1422–1434 (2013).

8. van Bergeijk, P., Adrian, M., Hoogenraad, C.C. & Kapitein, L.C. Optogenetic control of organelle transport and positioning. Nature 518, 111–114 (2015).

9. Gautier, A. et al. How to control proteins with light in living systems. Nat Chem Biol 10, 533–541 (2014).

10. Wu, C. et al. Recent Advances in Magnetic-Nanomaterial-Based Mechanotransduction for Cell Fate Regulation. Adv Mater 30, e1705673 (2018).

11. Monzel, C., Vicario, C., Piehler, J., Coppey, M. & Dahan, M. Magnetic control of cellular processes using biofunctional nanoparticles. Chemical science 8, 7330–7338 (2017).

12. Huang, H., Delikanli, S., Zeng, H., Ferkey, D.M. & Pralle, A. Remote control of ion channels and neurons through magnetic-field heating of nanoparticles. Nat Nanotechnol 5, 602–606 (2010).

13. Stanley, S.A. et al. Radio-wave heating of iron oxide nanoparticles can regulate plasma glucose in mice. Science 336, 604–608 (2012).

14. Etoc, F. et al. Subcellular control of Rac-GTPase signalling by magnetogenetic manipulation inside living cells. Nat Nanotechnol 8, 193–198 (2013).

15. Riggio, C. et al. The orientation of the neuronal growth process can be directed via magnetic nanoparticles under an applied magnetic field. Nanomedicine : nanotechnology, biology, and medicine 10, 1549–1558 (2014).

16. Stanley, S.A., Sauer, J., Kane, R.S., Dordick, J.S. & Friedman, J.M. Remote regulation of glucose homeostasis in mice using genetically encoded nanoparticles. Nature medicine 21, 92–98 (2015).

17. Wheeler, M.A. et al. Genetically targeted magnetic control of the nervous system. Nat Neurosci 19, 756–761 (2016).

18. Munshi, R. et al. Magnetothermal genetic deep brain stimulation of motor behaviors in awake, freely moving mice. eLife 6 (2017).

19. Schoneborn, H. et al. Novel Tools towards Magnetic Guidance of Neurite Growth: (I) Guidance of Magnetic Nanoparticles into Neurite Extensions of Induced Human Neurons and In Vitro Functionalization with RAS Regulating Proteins. J Funct Biomater 10 (2019).

20. Li, J.H. et al. Directed manipulation of membrane proteins by fluorescent magnetic nanoparticles. Nature communications 11, 4259 (2020).

21. Bongaerts, M. et al. Parallelized Manipulation of Adherent Living Cells by Magnetic Nanoparticles-Mediated Forces. International journal of molecular sciences 21 (2020).

22. Raudzus, F. et al. Magnetic spatiotemporal control of SOS1 coupled nanoparticles for guided neurite growth in dopaminergic single cells. Scientific reports 10, 22452 (2020).

23. Keizer, V.I.P. et al. Live-cell micromanipulation of a genomic locus reveals interphase chromatin mechanics. Science 377, 489–495 (2022).

24. Meister, M. Physical limits to magnetogenetics. eLife 5 (2016).

25. Nel, A., Xia, T., Madler, L. & Li, N. Toxic potential of materials at the nanolevel. Science 311, 622–627 (2006).

26. Nel, A.E. et al. Understanding biophysicochemical interactions at the nano-bio interface. Nat Mater 8, 543–557 (2009).

27. Lisse, D. et al. Monofunctional stealth nanoparticle for unbiased single molecule tracking inside living cells. Nano Lett 14, 2189–2195 (2014).

28. Verkman, A.S. Solute and macromolecule diffusion in cellular aqueous compartments. Trends Biochem Sci 27, 27–33 (2002).

29. Etoc, F. et al. Non-specific interactions govern cytosolic diffusion of nanosized objects in mammalian cells. Nature materials 17, 740–746 (2018).

30. Arakaki, A., Webb, J. & Matsunaga, T. A novel protein tightly bound to bacterial magnetic particles in Magnetospirillum magneticum strain AMB-1. J Biol Chem 278, 8745–8750 (2003).

31. Amemiya, Y., Arakaki, A., Staniland, S.S., Tanaka, T. & Matsunaga, T. Controlled formation of magnetite crystal by partial oxidation of ferrous hydroxide in the presence of recombinant magnetotactic bacterial protein Mms6. Biomaterials 28, 5381–5389 (2007).

32. Wang, L. et al. Self-assembly and biphasic iron-binding characteristics of Mms6, a bacterial protein that promotes the formation of superparamagnetic magnetite nanoparticles of uniform size and shape. Biomacromolecules 13, 98–105 (2012).

33. Shipunova, V.O. et al. Self-assembling nanoparticles biofunctionalized with magnetite-binding protein for the targeted delivery to HER2/neu overexpressing cancer cells. J Magn Magn Mater 469, 450–455 (2019).

34. Rawlings, A.E., Liravi, P., Corbett, S., Holehouse, A.S. & Staniland, S.S. Investigating the ferric ion binding site of magnetite biomineralisation protein Mms6. PLoS One 15, e0228708 (2020).

35. Yamagishi, A., Narumiya, K., Tanaka, M., Matsunaga, T. & Arakaki, A. Core Amino Acid Residues in the Morphology-Regulating Protein, Mms6, for Intracellular Magnetite Biomineralization. Scientific reports 6, 35670 (2016).

36. Kirchhofer, A. et al. Modulation of protein properties in living cells using nanobodies. Nat Struct Mol Biol 17, 133–138 (2010).

37. Lisse, D. et al. Engineered Ferritin for Magnetogenetic Manipulation of Proteins and Organelles Inside Living Cells. Advanced materials 29, 1700189 (2017).

38. Machacek, M. et al. Coordination of Rho GTPase activities during cell protrusion. Nature 461, 99–103 (2009).

39. Pertz, O. Spatio-temporal Rho GTPase signaling - where are we now? J Cell Sci 123, 1841–1850 (2010).

40. Nobes, C.D. & Hall, A. Rho GTPases control polarity, protrusion, and adhesion during cell movement. J Cell Biol 144, 1235–1244 (1999).

41. Mehidi, A. et al. Transient Activations of Rac1 at the Lamellipodium Tip Trigger Membrane Protrusion. Curr Biol 29, 2852–2866 e2855 (2019).

42. Wu, Y.I. et al. A genetically encoded photoactivatable Rac controls the motility of living cells. Nature 461, 104–108 (2009).

43. Lafontaine, D.L.J., Riback, J.A., Bascetin, R. & Brangwynne, C.P. The nucleolus as a multiphase liquid condensate. Nat Rev Mol Cell Biol (2020).

44. Hildebrand, E.M. & Dekker, J. Mechanisms and Functions of Chromosome Compartmentalization. Trends in biochemical sciences 45, 385–396 (2020).

45. Rhine, K., Vidaurre, V. & Myong, S. RNA Droplets. Annual review of biophysics 49, 247–265 (2020).

46. Ong, J.Y. & Torres, J.Z. Phase Separation in Cell Division. Mol Cell (2020).

47. Wu, X., Cai, Q., Feng, Z. & Zhang, M. Liquid-Liquid Phase Separation in Neuronal Development and Synaptic Signaling. Dev Cell (2020).

48. Case, L.B., Ditlev, J.A. & Rosen, M.K. Regulation of Transmembrane Signaling by Phase Separation. Annu Rev Biophys 48, 465–494 (2019).

49. Nozawa, R.S. et al. Nuclear microenvironment in cancer: Control through liquid-liquid phase separation. Cancer science 111, 3155–3163 (2020).

50. Alberti, S. & Dormann, D. Liquid-Liquid Phase Separation in Disease. Annual review of genetics 53, 171–194 (2019).

51. Shin, Y. & Brangwynne, C.P. Liquid phase condensation in cell physiology and disease. Science 357 (2017).

52. Nott, T.J. et al. Phase transition of a disordered nuage protein generates environmentally responsive membraneless organelles. Mol Cell 57, 936–947 (2015).

53. Shin, Y. et al. Spatiotemporal Control of Intracellular Phase Transitions Using Light-Activated optoDroplets. Cell 168, 159–171 e114 (2017).

54. Capecchi, M.R. High efficiency transformation by direct microinjection of DNA into cultured mammalian cells. Cell 22, 479–488 (1980).

55. Gavutis, M., Lata, S., Lamken, P., Müller, P. & Piehler, J. Lateral ligand-receptor interactions on membranes probed by simultaneous fluorescence-interference detection. Biophys J 88, 4289–4302 (2005).

56. Wedeking, T. et al. Spatiotemporally Controlled Reorganization of Signaling Complexes in the Plasma Membrane of Living Cells. Small 11, 5912–5918 (2015).

57. Rawlings, A.E. et al. Ferrous Iron Binding Key to Mms6 Magnetite Biomineralisation: A Mechanistic Study to Understand Magnetite Formation Using pH Titration and NMR Spectroscopy. Chemistry 22, 7885–7894 (2016).

