## Supplemental Figures for "Biofunctional coating of synthetic magnetic nanoparticles enables magnetogenetic control of protein functions inside cells"

### Supplementary Figures

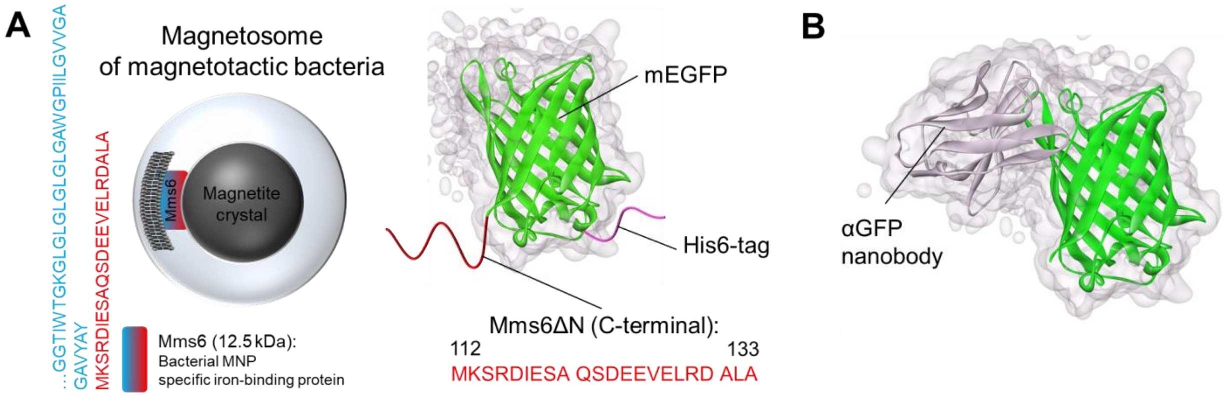


**Figure S1** **Genetic engineering of mEGFP for MCP biofunctionalization.** (A) Mms6 – bacterial magnetic particle specific iron-binding protein is part of the magnetosome membrane of magnetotactic bacteria (adapted from ref[57]). (B) Biofunctionalization of mEGFP-coated MCP via the αGFP – mEGFP interaction (pdb: 3K1K).

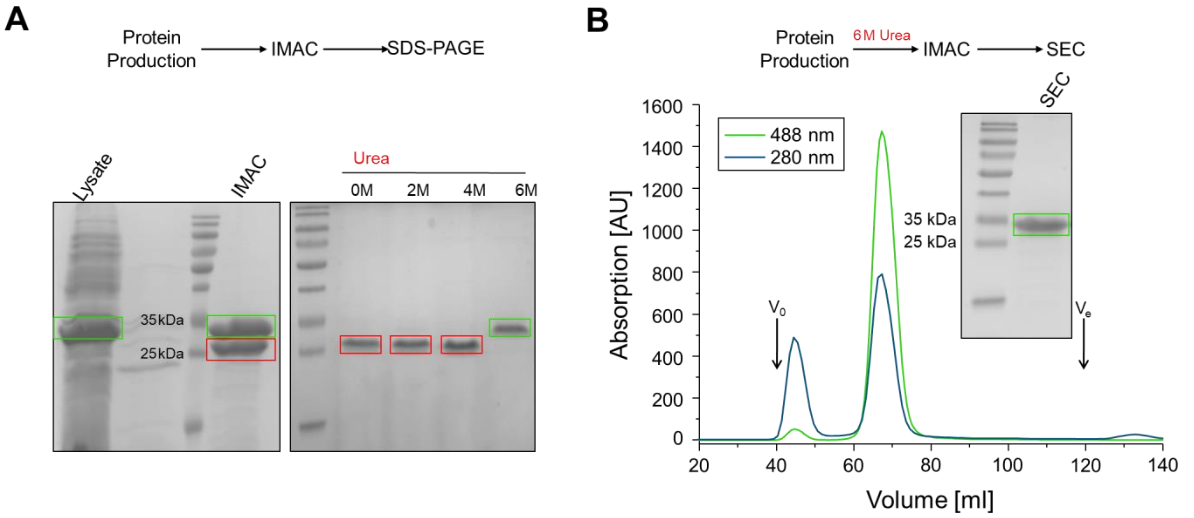


**Figure S2** **Purification of H6::mEGFP::Mms6.** (A) Optimization of a standard protocol for purification of H6::mEGFP::Mms6. Left: SDS-PAGE after immobilized metal affinity chromatography (IMAC) of H6::mEGFP::Mms6 according to a standard protocol for HIS-Tagged proteins. Full length protein is highlighted by the green rectangle and proteolytic products by the red rectangle. Right: optimization of the purification strategy by titration of different urea concentrations. Cells were lysed in presence of urea. (B) Final purification strategy in presence of 6M urea. Size-exclusion chromatography (SEC) and SDS-PAGE of H6::mEGFP::Mms6 after SEC.

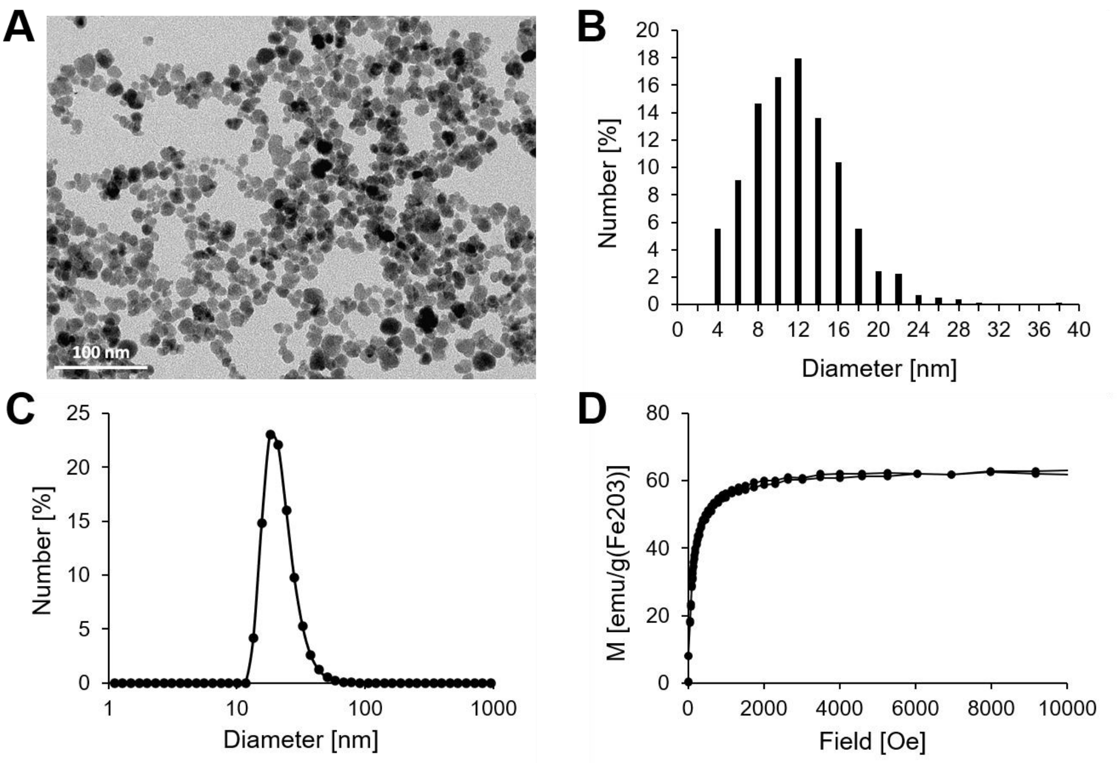


**Figure S3** **Characterization of the maghemite MCP.** (A) Transmission electron microscopy image of MCPs. (B) Physical size distribution of the MCPs determined by measuring over 1250 particles with ImageJ yielded a mean diameter of 11.7 nm. (C) Hydrodynamic diameter distribution of the MCPs in water at pH = 2. (D) Magnetization curve of the MCP dispersed in water at pH = 2. The saturation magnetization is 62.5 emu/g($Fe_2O_3$).

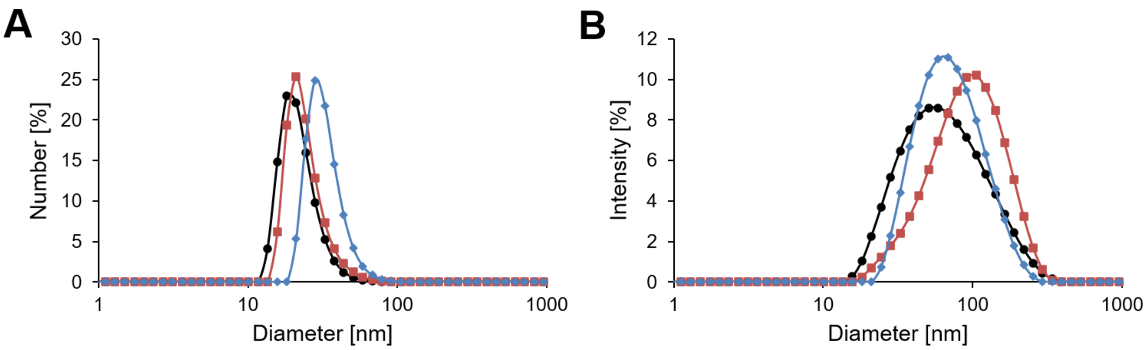


**Figure S4** **Hydrodynamic diameter analysis of the particles.** Size distributions (A) in number and B) in intensity obtained by dynamic light scattering of bare MCP (black circles), MCP functionalized with non-PEGylated Mms6-GFP (orange squares) and MCP functionalized with PEGylated Mms6-GFP (light blue diamonds). An increase in the hydrodynamic diameter of the MCP is observed with both proteins, with a larger increase in presence of the PEG chains when looking at the size distribution in number. Additionally, the effect of PEGylation on the colloidal stability of the syMagIcs can be inferred in intensity. The shift towards the larger diameter for the MCP functionalized with non-PEGylated Mms6-GFP compared to the MCP functionalized with PEGylated Mms6-GFP can be explained by some aggregation of the non-PEGylated particles.

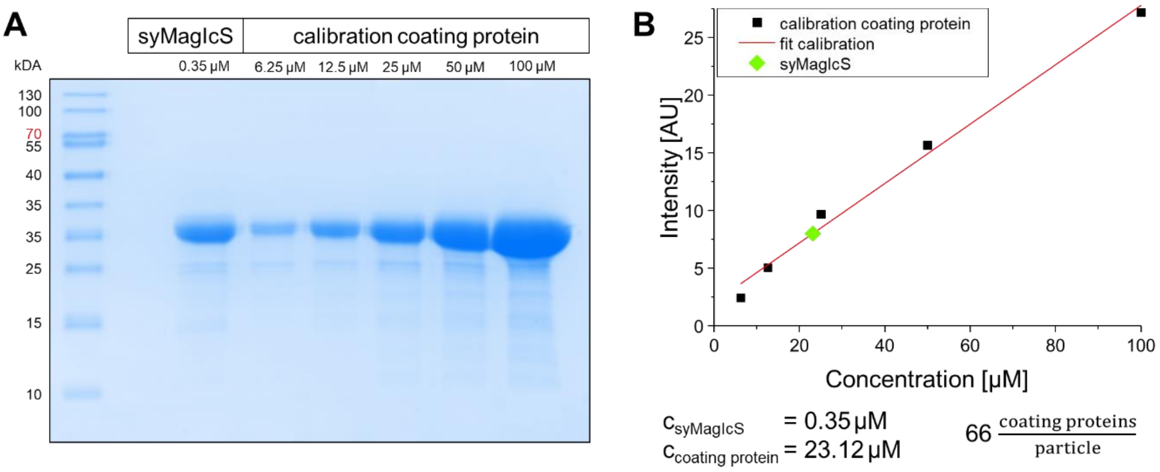


**Figure S5** **Estimation of functionalization stoichiometry.** (A) Denaturating SDS-PAGE of syMagIcS and different concentration of coating protein. Coomassie staining. (B) Intensity in dependence on coating protein concentration. Calculation of coating protein concentration bound to syMagIcS and functionalization stoichiometry.

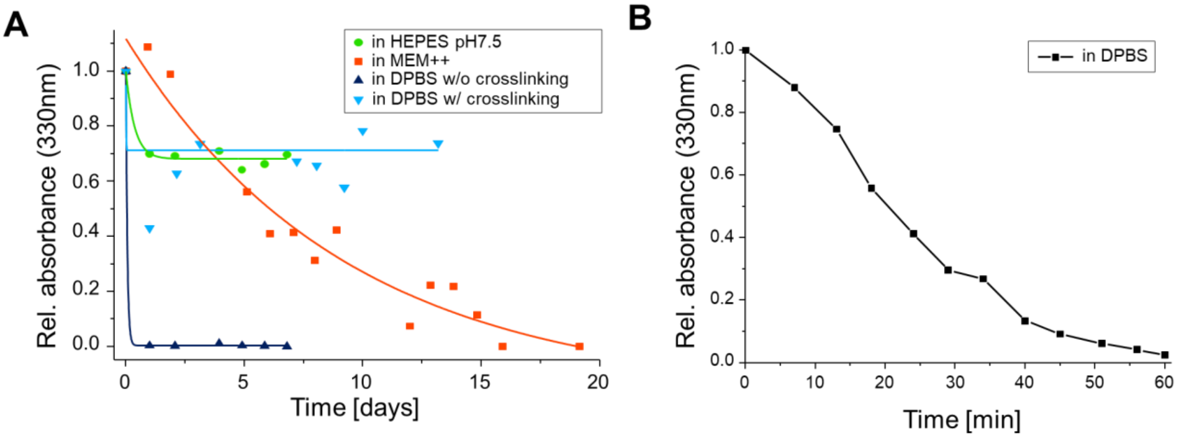


**Figure S6** **Long-term particle stability under different conditions.** (A) Stability of syMagIcS in different physiological solutions and crosslinking of the particle coating using 4% PFA. (B) Stability of syMagIcS in DPBS over 1 h.

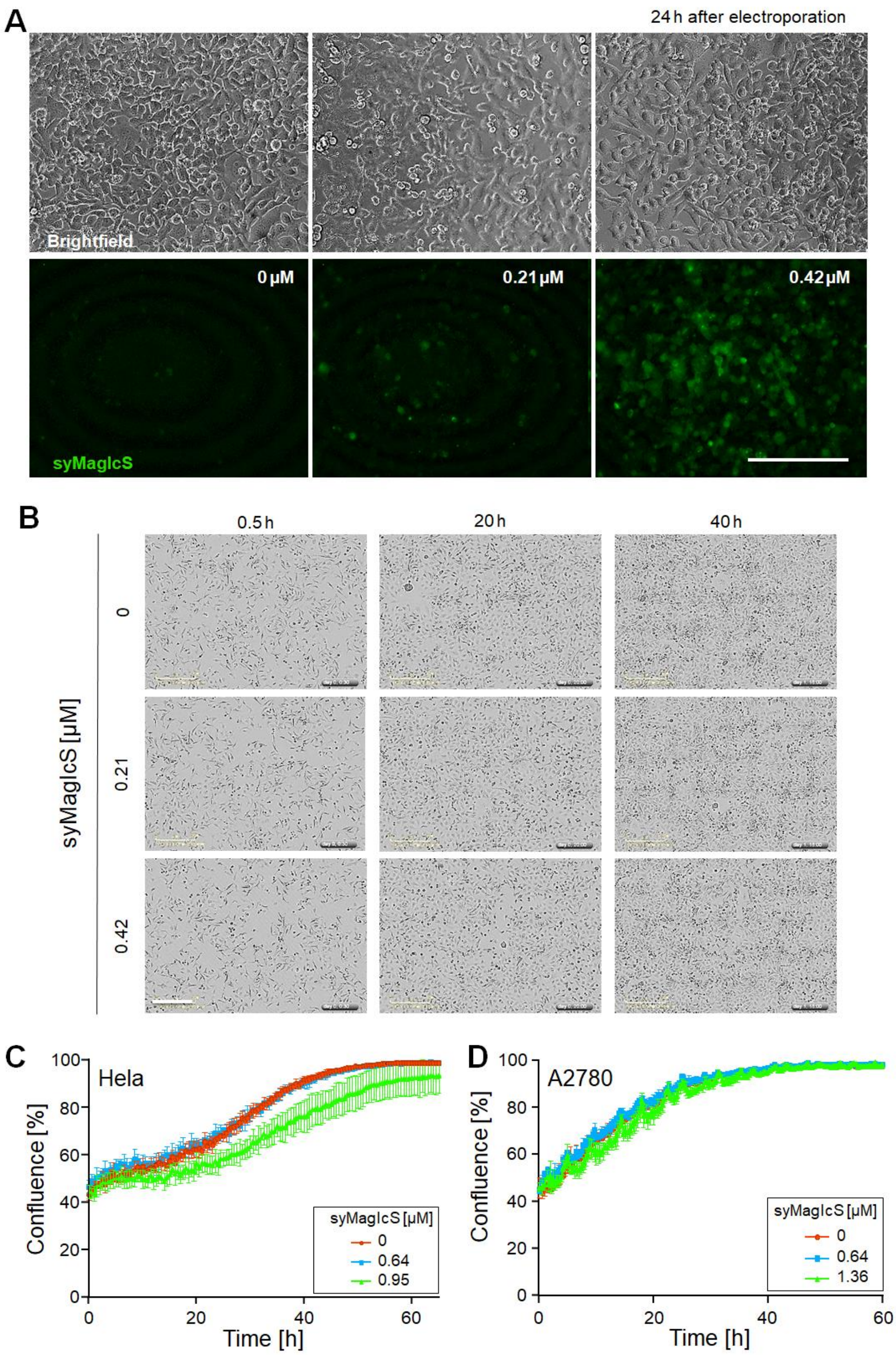


**Figure S7 Cytotoxicity of syMagIcS analysed by IncuCyte automated imaging system.** (A) Intracellular delivery of syMagIcS into HeLa cells using electroporation at different bulk concentrations. Images were taken 24 hours after electroporation using widefield fluorescence microscopy. Phase-contrast transmission (top) and back-ground-corrected green fluorescence (bottom) images for different syMagIcS bulk concentrations as indicated. Scale bar: 300 µm. (B) Representative images of Hela cells obtained by the IncuCyte system used for calculation of cell confluence. Scale bar: 300 µm. (C, D) Time-lapse cell confluence of Hela (C) and A2780 (D) cells that were electroporated with different concentration of syMagIcS. Time 0 h represents cells 4-5 hour after electroporation. Data points show the mean ± SD from 3 independent experiments.

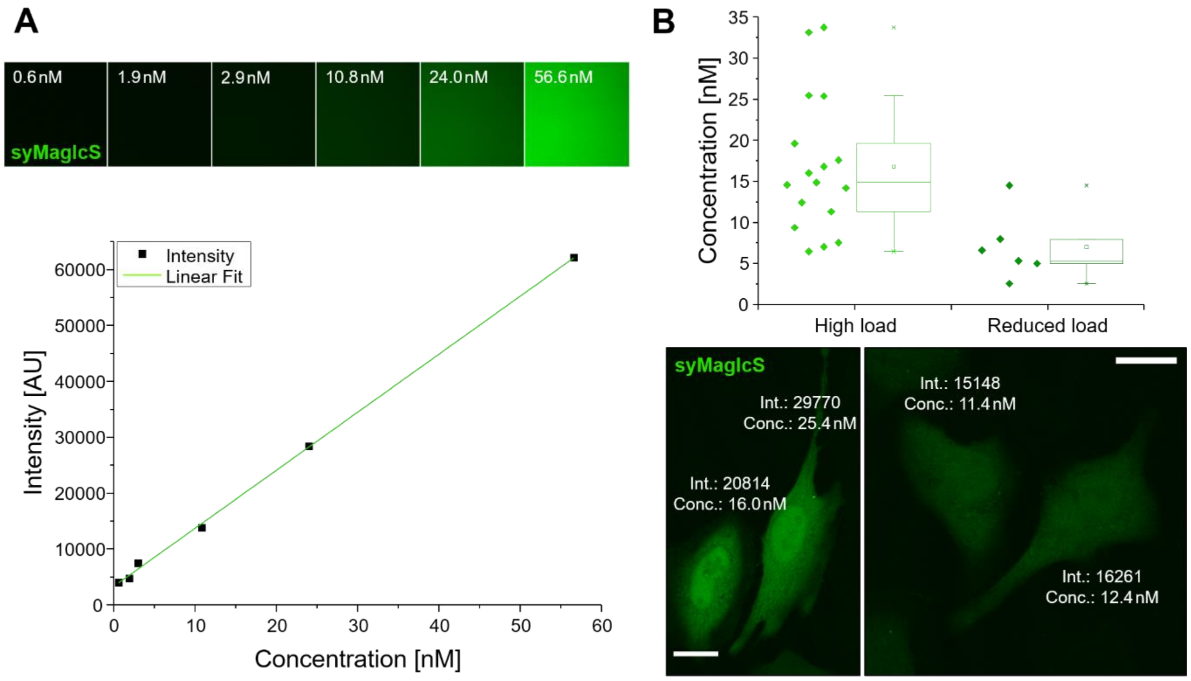


**Figure S8 Quantification of syMagIcS concentrations inside cells**. (A) Confocal laser scanning microscopy images of syMagIcS solutions at different concentrations. (B) Concentration measurements inside cells. Scale bars: 20 µm in all images.

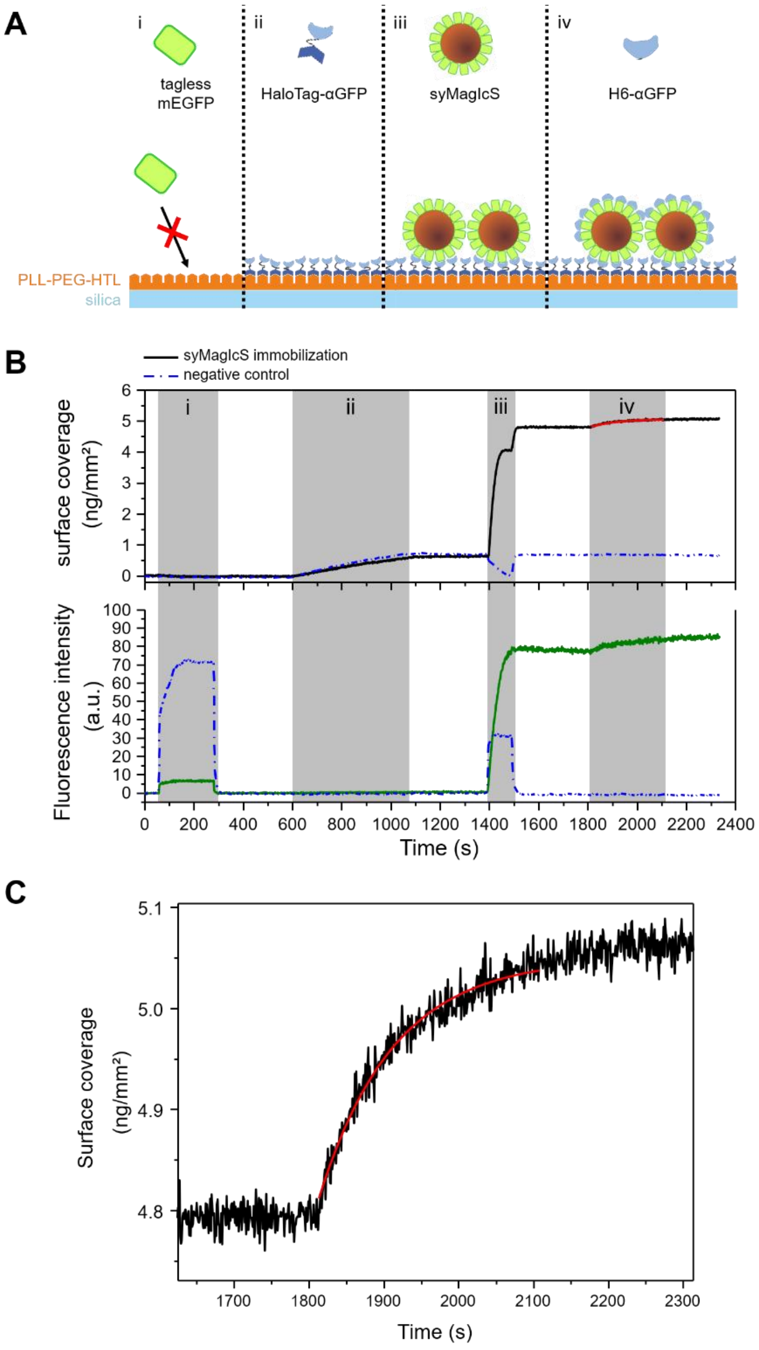


**Figure S9** **Interaction of αGFP with syMagIcS quantified by solid phase detection *in vitro*.** (A) Cartoon of the assay comprising: (i) injection of tagless mEGFP to ensure no nonspecific binding to the PLL-PEG-HTL surface, (ii) immobilization of HaloTag-H10-αGFP to HTL ligands at a concentration of 1μM; (iii) binding of syMagIcS to immobilized αGFP; (iv) binding of αGFP to toplayer of immobilized syMagIcS. (B) Mass signal (top) and fluorescence signal (bottom) acquired during the assay. (C) Zoom into the mass signal detected for αGFP binding to immobilized syMagIcS (iv) and fit of Langmuir model ($k_{on}$: (4.0 ±0.1)×10$^4$ M$^{-1}$s$^{-1}$).

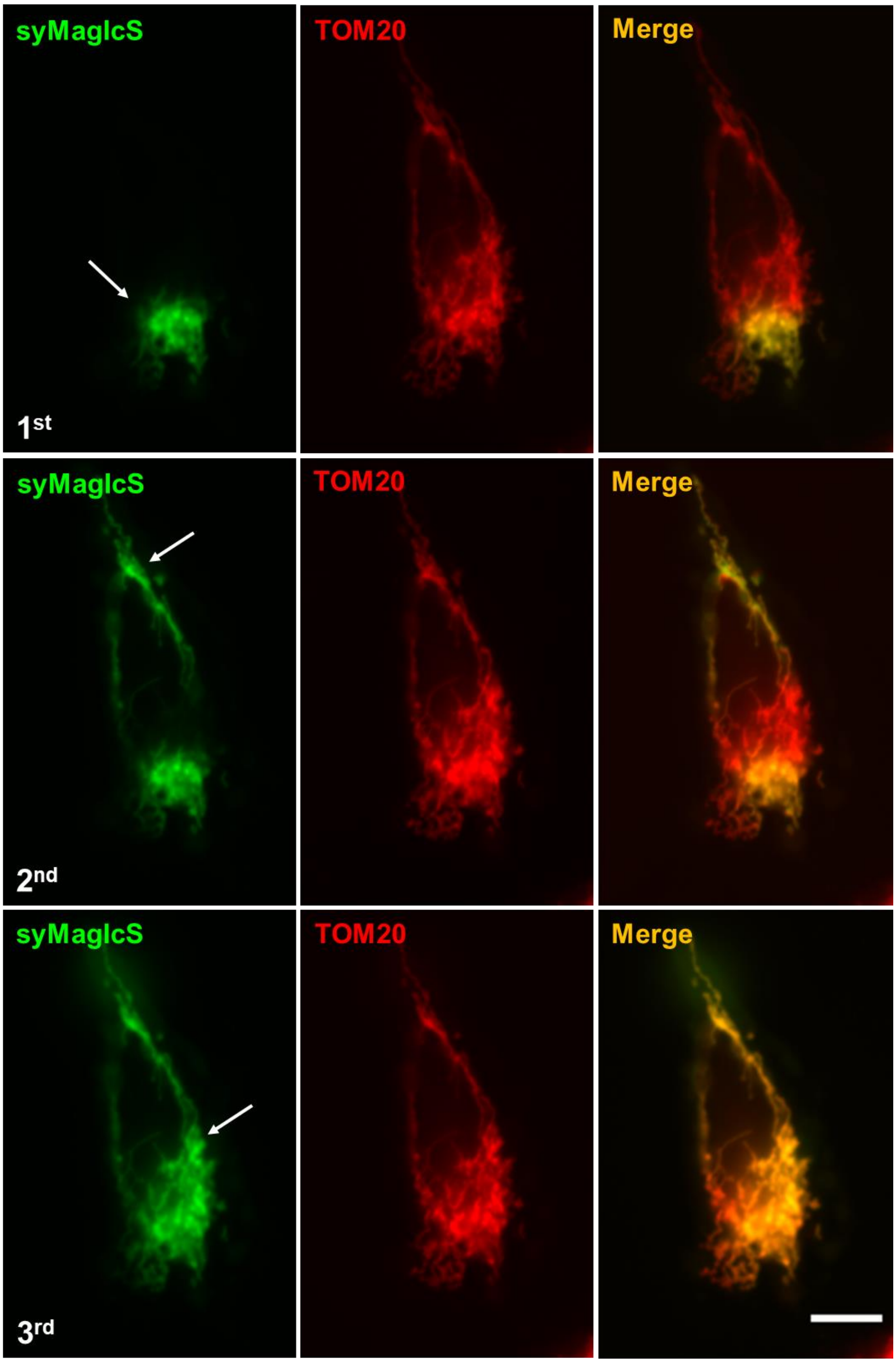


**Figure S10 Site-specific targeting to mitochondria.** Rapid site-specific targeting of syMagIcS (green) upon repeated micro-injection into HeLa cell transiently overexpressing TOM20::mCherry::αGFP (red). Scale bar: 10 µm.

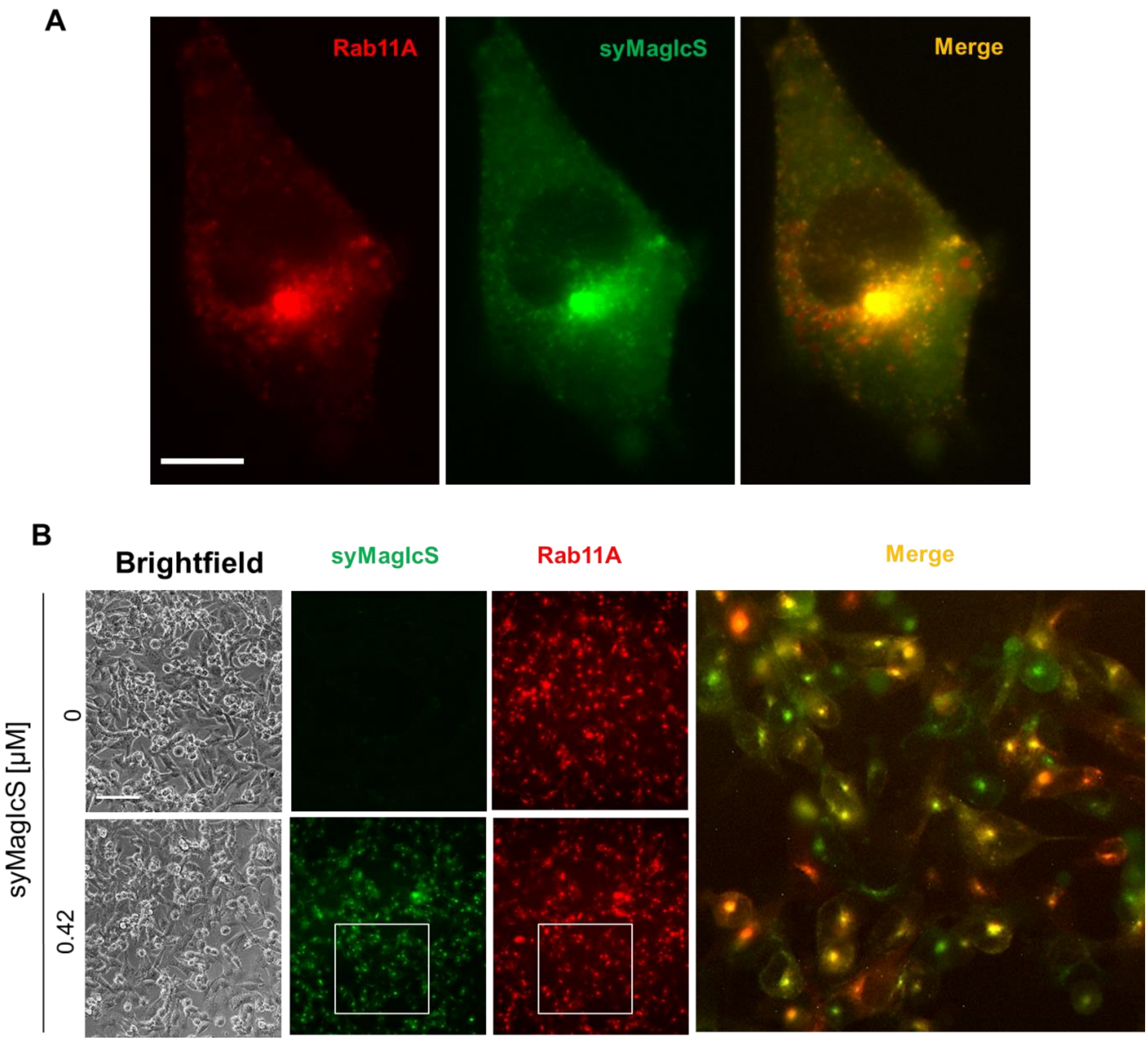


**Figure S11 Site-specific targeting of syMagIcS recycling endosomes inside living cells.** syMagIcS (green) in the cytosol of A2780 cells stably over-expressing αGFP::mCherry:Rab11A (red). (A) Single cell with syMagIcS (green) being microinjected into the cytoplasm. Scale bar: 30 µm (B) Images 24 h after being electroporated with 0 or 0.42 µM of syMagIcS. Highlighted area is enlarged with merged green and red signal. Background was subtracted using ImageJ in green images. Scale bar: 300 µm.

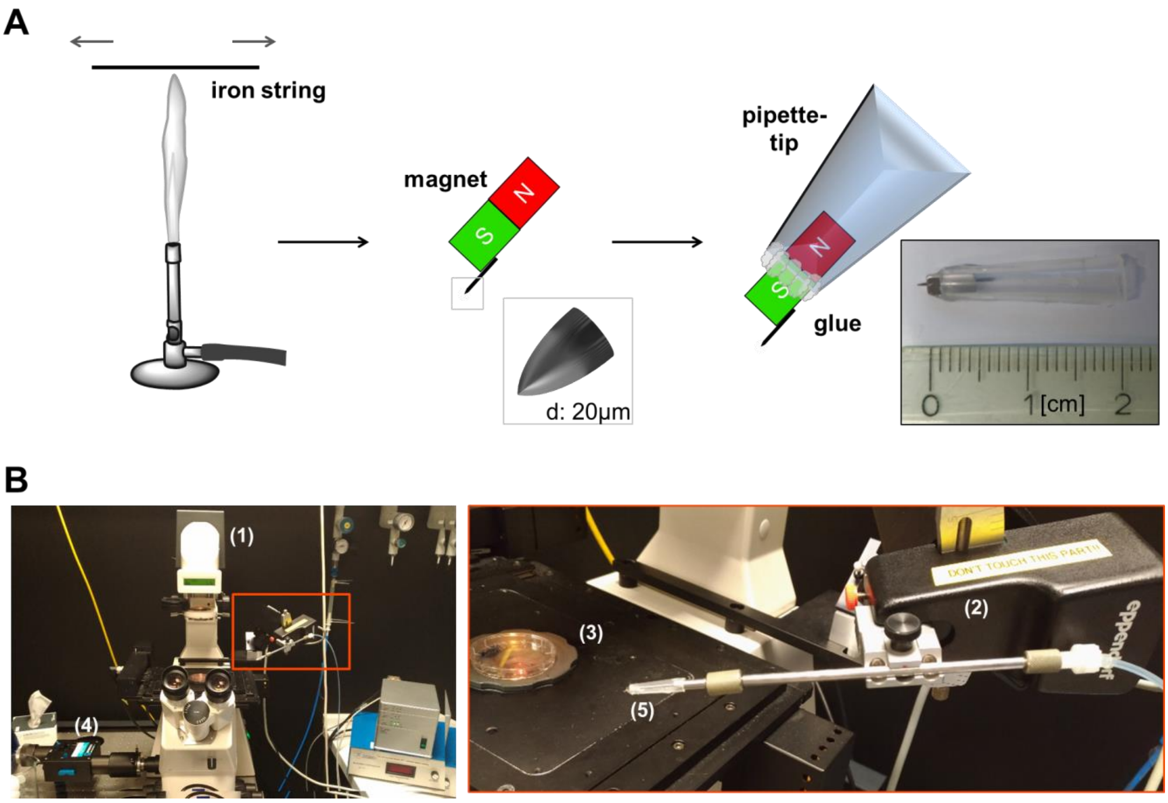


**Figure S12 Fabrication of the magnetic tip and its application for magnetic manipulation.** (A) Tip fabrication and assembly: An iron wire is pulled over a Bunsen burner flame to yield a fine tip, which is attached to a magnet (Ni-45SH) and glued into a plastic pipette tip. This allows the magnetic tip to be attached to the micromanipulation apparatus. (B) Images of the microscope setup for the magnetic manipulation of human cells. (1) Microscope stand, (2) Micromanipulator, (3) Cell sample, (4) Camera and (5) Micromagnet attached to micromanipulator.

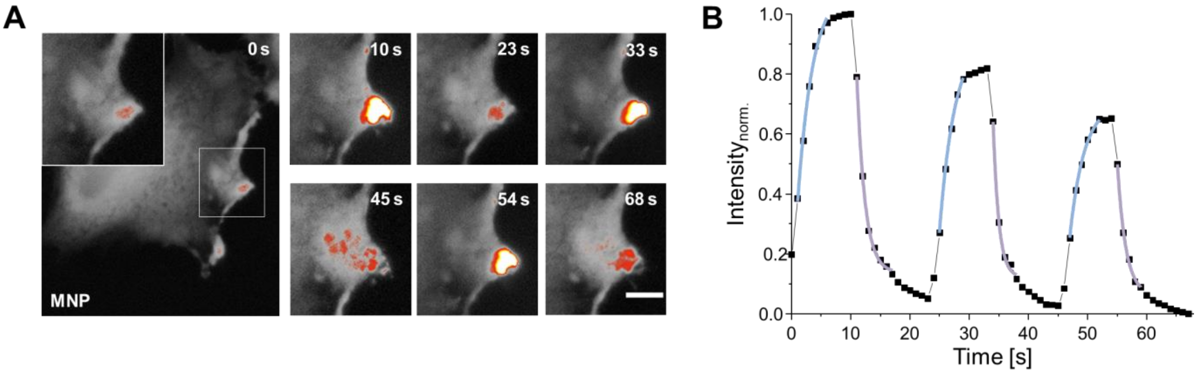


**Figure S13 syMagIcS attraction and release kinetics in living cells.** (A) syMagIcS were injected into COS7 cells and repeatedly attracted and released with a micro-magnet to the plasma membrane. Scale bar: 10 µm. (B) Attraction and release kinetics of syMagIcS determined from the changes in fluorescence intensity in the area highlighted by the white square and exponential fit of the curve (blue, purple).

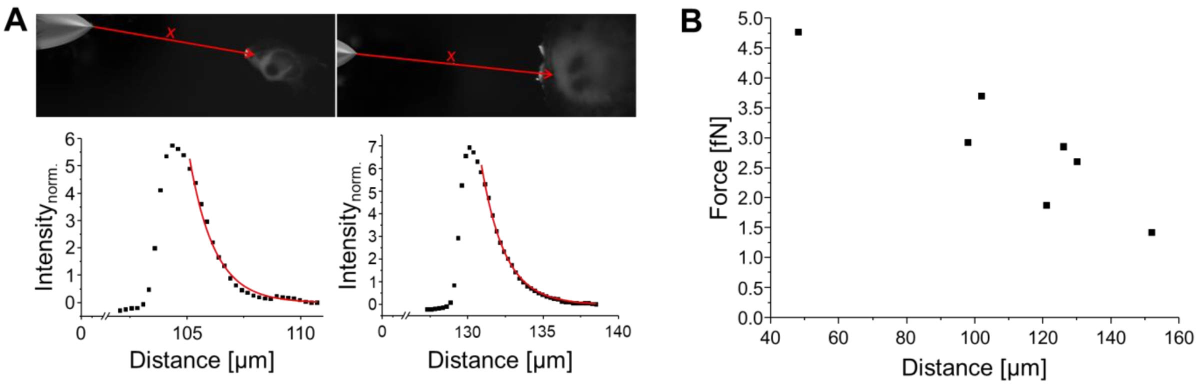


**Figure S14 Estimation of force exerted from particles.** (A) Microscopy images and steady-state syMagIcS gradient profiles inside the cytoplasm of living cells generated by a magnetic gradient. The decay into the cytoplasm was fitted by an exponential decay to estimate the force exerted on the MNPs (SI Methods, Equation 1). (B) Force observed for the magnetic tip places at different distances from the cell.

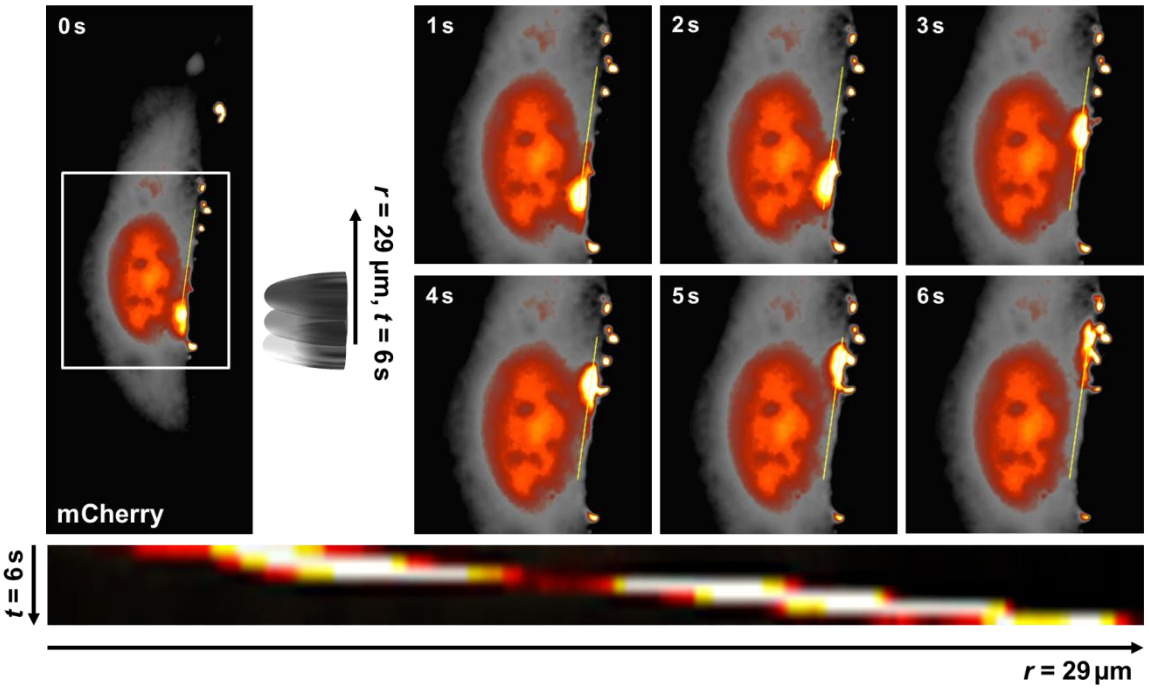


**Figure S15 Rapid relocalization of proteins by magnetic manipulation.** syMagIcS were injected into a HeLa cell expressing αGFP::mCherry (red hot). After approaching the magnetic tip, its position was continuously shifted within a total time interval for 6 s and the reorganization of mCherry was followed by time-lapse microscopy.

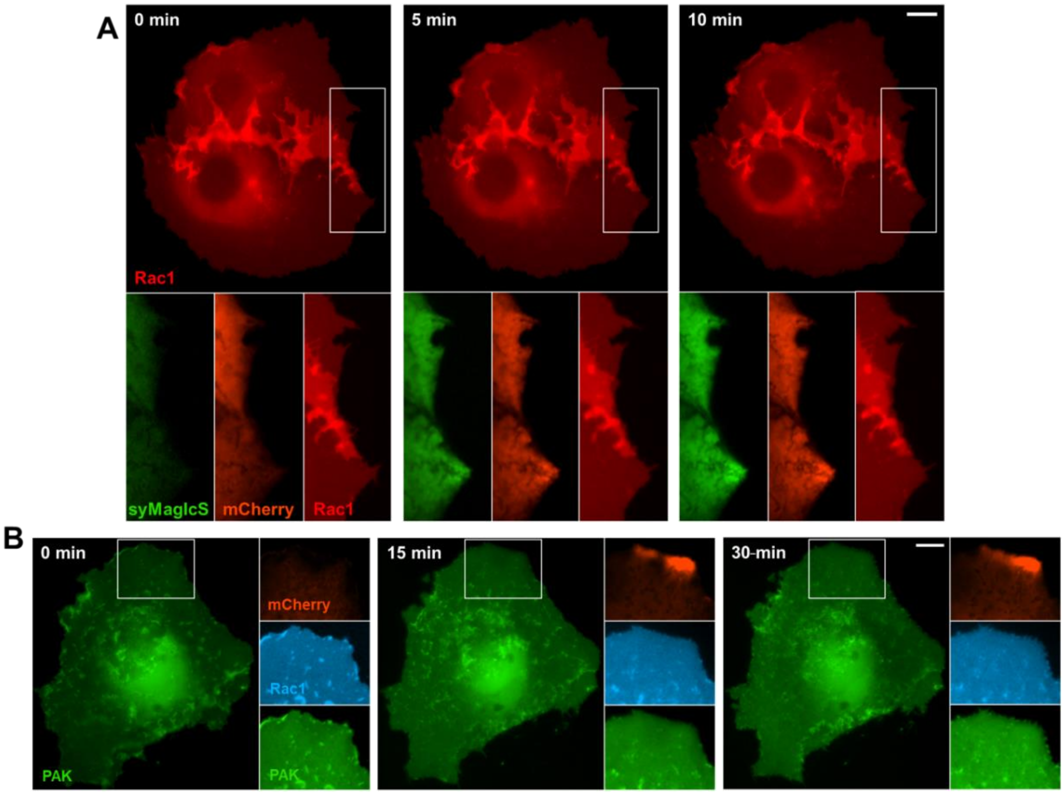


**Figure S16 Negative control experiments of magnetogenetic Rac1 activation.** (A) Negative control of Rac1 recruitment. A COS7 cell expressing SiR-labeled HaloTag::Rac1 (red) and αGFP::mCherry (orange) after injection of syMagIcS (green). Overview image ($^{SIR}$HaloTag::Rac1) and crop of syMagIcS (green), mCherry (orange) and Rac1 (red) at different time points during magnetic manipulation. (B) Negative control of magnetogenetic Rac1 activation. COS7 cell expressing αGFP::mCherry (orange) and mNeonGreen::PAK-CRIB::P2A::Rac1::mTFP1::CAAX (green/cyan) after injection of non-fluorescent syMagIcS. Overview image showing PAK and crop at the ROI indicated by a white rectangle for all three channels at different time points after initiating magnetic manipulation. Scale bar: 10 µm.

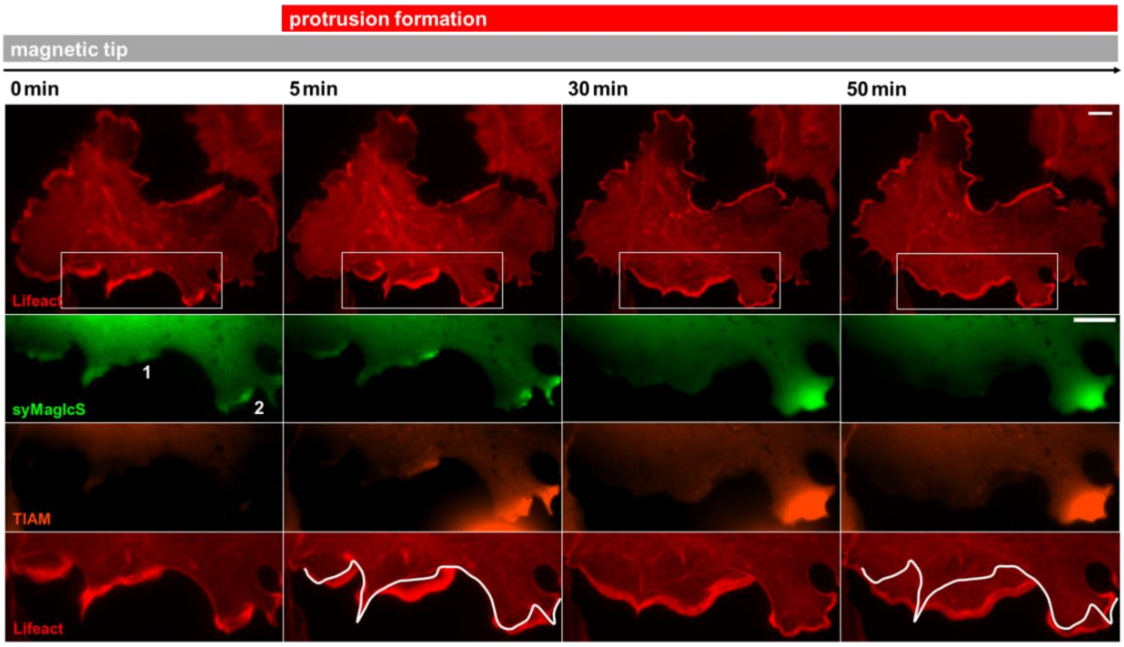


**Figure S17 Magnetogenetic activation of Rac1.** syMagIcS (green) was injected into cells expressing TIAM1-DHPH::mCherry::αGFP (orange) and Lifeact::iRFP (red, overview and crop) for staining f-actin. Time-lapse imaging during magnetic manipulation showing the actin cytoskeletal structure in regions with low (1) and high (2) syMagIcS densities. Scale bar: 10 µm.

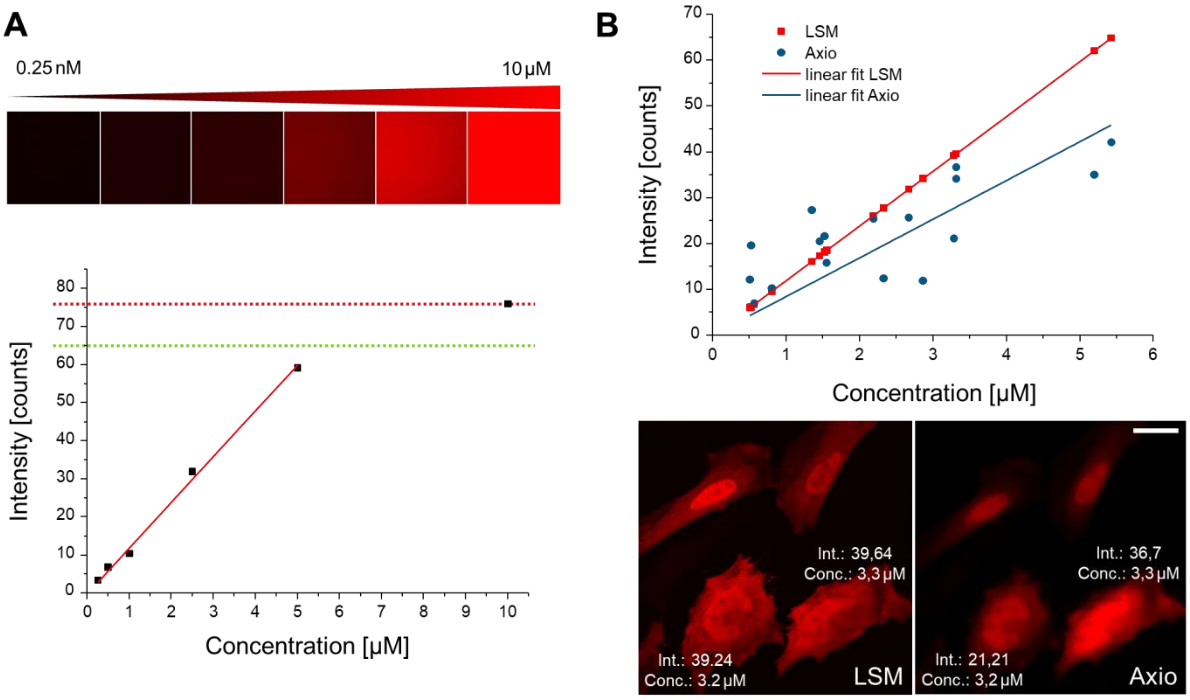


**Figure S18: Determination of the DDX4 concentration in the cytoplasm.** (A) Calibration using differently concentrated DDX4::mCherry::αGFP solutions. (B) Concentration measurements inside cells. Scale bar: 20 μm.

### Supplementary Movies

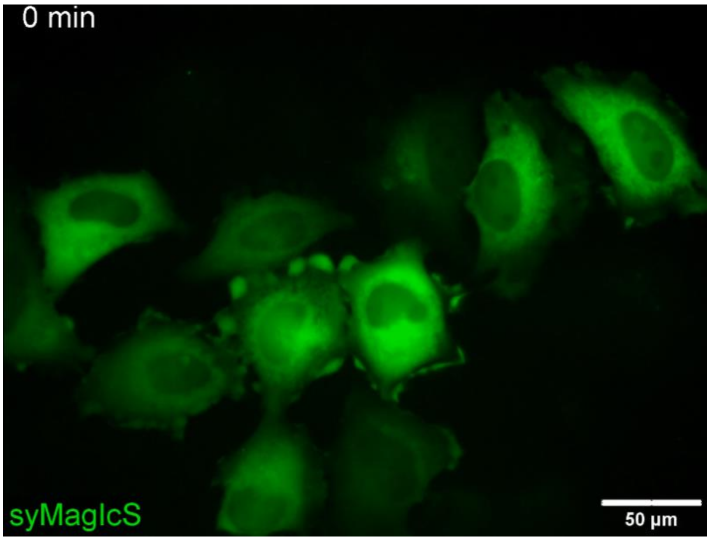


Movie S1 Homogeneous dispersion of syMagIcS in the cytosplasm of HeLa cells.

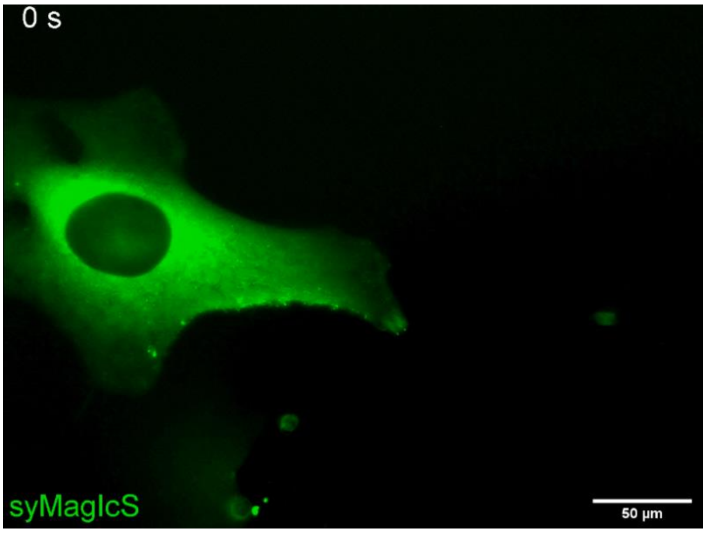


Movie S2 Manipulation syMagIcS by a magnetic field gradient.

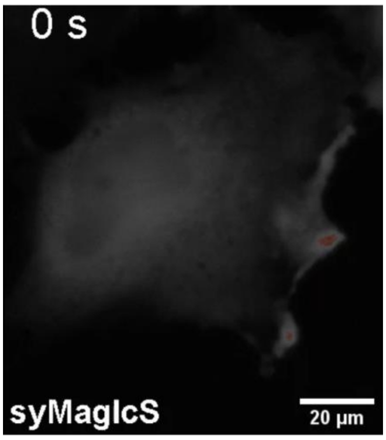


Movie S3 Reversible attraction of syMagIcS.

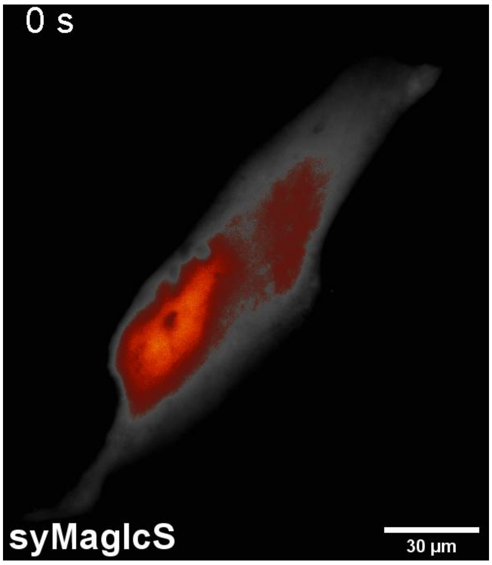


Movie S4 Magnetogenetic manipulation of mCherry::αGFP captured to syMagIcS.

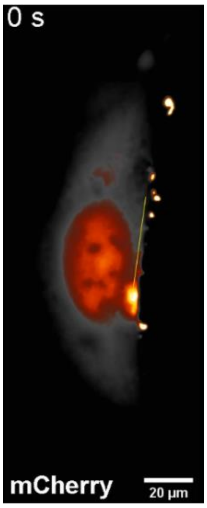


Movie S5 Fast re-localization of mCherry in the cytoplasm of HeLa cells.

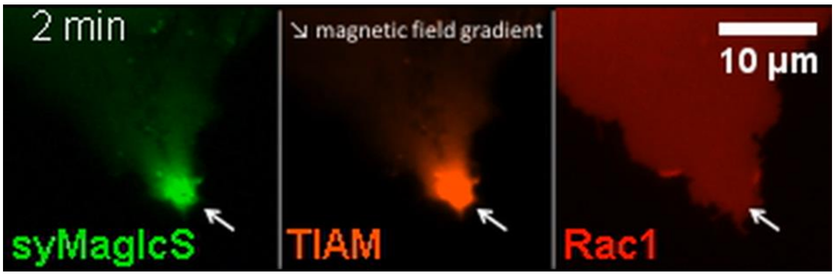


Movie S6 Recruitment of Rac1 controlled by TIAM-functionalized syMagIcS.

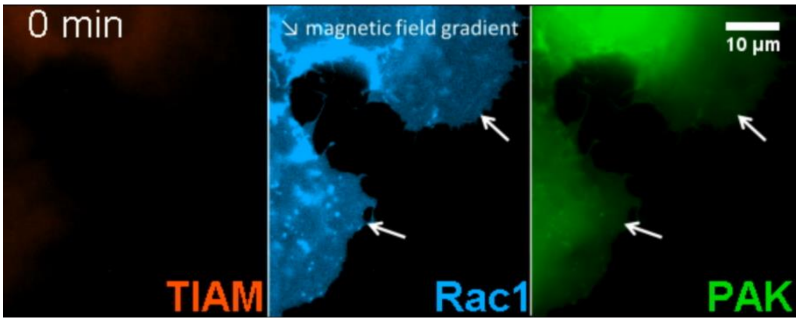


Movie S7 Magnetogenetic activation of Rac1 probed by PAK-CRIB recruitment.

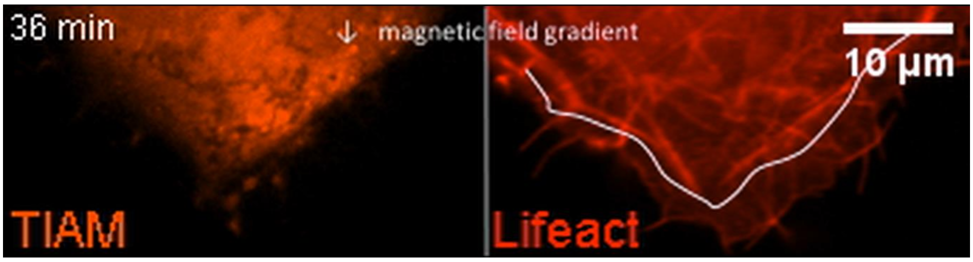


Movie S8 Magnetogenetic activation of Rac1 downstream signalling.

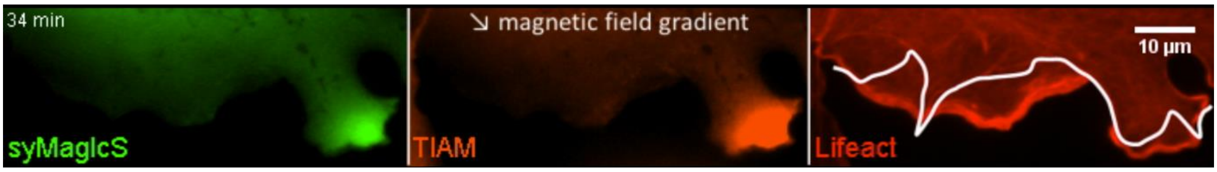


Movie S9 Magnetogenetic activation of Rac1 downstream signalling.

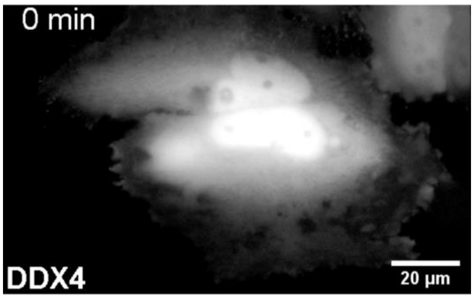


Movie S10 Spatial control of DDX4 LLPS via magnetic mECFP-coated syMagIcS.
